# The Earliest Impressions: A Systematic Review of Early-Life Exposures on Brain Structure and Neurodevelopmental Outcomes

**DOI:** 10.64898/2026.07.09.737480

**Authors:** Judith Dehnen, Harvey Brown, Aaron Alexander-Bloch, Richard A. I. Bethlehem

## Abstract

Prenatal and early postnatal life is a period of rapid brain growth, making the developing brain particularly susceptible to external influences. Adapting the biopsychosocial model of mental health and illness, this review provides a systematic overview of how biological, psychological, and social exposures from conception to age three critically converge to shape brain development and neurodevelopmental outcomes. Following a pre-registered protocol and the Preferred Reporting Items for Systematic Reviews and Meta-Analysis (PRISMA) guidelines, 55 studies were included, primarily published in the past 15 years. Earlier studies focused predominantly on biological exposures, while more recent work has increasingly examined psychological exposures and, more rarely, social exposures. While each exposure exhibited its own pattern of brain alterations and neurodevelopmental changes, an overarching pattern emerged across the different components of the biopsychosocial model. Adverse biological exposures were consistently associated with delayed brain maturation as reflected by brain imaging measures. Adverse psychosocial exposures showed a more complex pattern of associations with both delayed and accelerated brain maturation. Crucially, adverse exposures, whether associated with delayed or accelerated brain maturation, were consistently associated with poorer neurodevelopmental outcomes, underscoring the necessity of considering both brain and behavior when estimating the impact of early exposures. We conclude that research into early-life exposures on brain maturation and neurodevelopmental outcomes is on the rise, but there is a great need for further investigation, in particular of psychological and social exposures. The interactions between exposures, the brain, and outcomes are highly complex, requiring assessment of both brain development and behavior together, ideally in within-subject longitudinal designs in future studies.

## Introduction

During the prenatal and early postnatal period, the brain develops at a pace unparalleled at any other point in life (1,2). During early pregnancy, most neurons are formed, migrate to their final position, and begin forming connections (2–4). In the last weeks of pregnancy, exuberant synaptogenesis begins, leading to a peak of synaptic connections around year 2, with pruning refining this architecture in parallel (4,5). Simultaneously, myelin begins forming around neurons in the third trimester, a process largely complete by the second postnatal year but continuing into adulthood in some regions (2,6). As a result, brain volume reaches roughly 80% of adult size by age 2, alongside rapid increases in cortical thickness, surface area, and gyrification (1–3).

These early neural changes are adaptive but make the developing brain particularly vulnerable to adverse exposures (2,7–9). Altered brain development has in turn been hypothesized to alter neurodevelopmental and behavioral outcomes (10,11). Yet, the relationship between biopsychosocial exposures, brain development, and behavioral outcomes is complex and understudied (7). To estimate the impact of different exposures on development, the full pathway must be evaluated within the same population: biopsychosocial exposures, brain development, and neurodevelopmental outcomes (see Figure 1 for a schematic overview).

**Figure 1:**
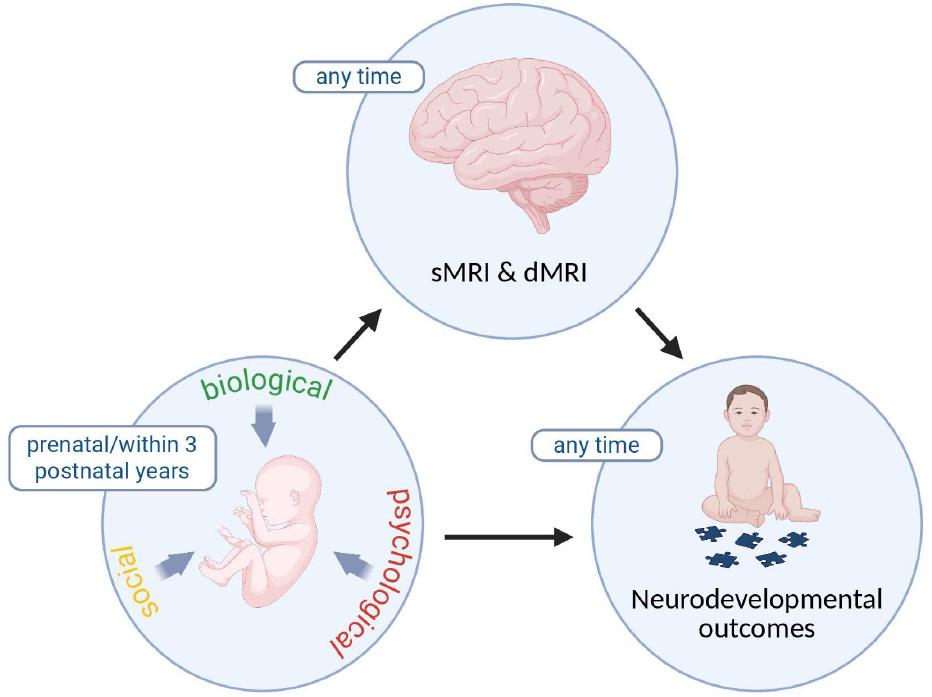
Proposed model of the exposure-brain-outcome pathway. The model illustrates the interplay between early-life exposures, brain development, and neurodevelopmental outcomes. Eligible timeframes and MRI modalities used as inclusion criteria in this review are indicated. sMRI, structural magnetic resonance imaging; dMRI, diffusion weighted magnetic resonance imaging. This figure was created in BioRender, https://BioRender.com.

Previous reviews have addressed these domains mainly in pairs: how specific exposures shape brain imaging measures (10–18), how exposures affect neurodevelopmental outcomes (19–23), or how brain alterations relate to neurodevelopmental outcomes (24–28). A few reviews examined the exposure-brain-outcome pathway for single exposures, but often relied on vague proxies for brain development (e.g., head circumference) or treated neuroimaging peripherally (29–36).

Therefore, this review provides a holistic overview of various pre- and postnatal exposures that shape the brain and neurodevelopmental outcomes. We systematically reviewed studies that assessed a prenatal or early postnatal exposure (until the age of 3 years), a measurement of brain morphology at any point in life, and any neurodevelopmental outcome. Given the focus on exposures rather than intrinsic characteristics of the developing child, this review does not include genetic factors. The exposures identified in this review are grouped following the biopsychosocial model (37) – a framework positing that biological, psychological, and social factors interact to determine health and developmental vulnerability. These exposures are briefly introduced below; more details can be found in the Supplementary Information.

### Biological exposures

Maternal immune activation during pregnancy may disrupt neurodevelopment, e.g., through raised inflammatory markers and pathogens crossing the placenta (38). Fetal growth restriction, often driven by placental insufficiency, limits oxygen and nutrient delivery to the developing brain (39,40). Prenatal drug exposure can alter neurotransmitter systems, particularly the dopaminergic system (35). Environmental toxicants such as air pollution, perfluoroalkyl substances (PFAS), and pesticides may alter brain development, e.g., through neurotransmitter disruption (41,42). Prematurity (preterm birth/very low birth weight (VLBW)) exposes the brain to the extrauterine environment early, often causing hypoxia-ischemia and inflammation (32,43), but even variation within the normal gestational range may subtly affect brain development (44). Neonatal complications can affect the brain, commonly via inflammation and hypoxia (45–47). Finally, early-life nutrition supports energy-demanding processes like myelination and neurogenesis (10).

### Psychological exposures

Maternal distress can affect the fetal brain, e.g., through Hypothalamic-Pituitary-Adrenal (HPA) axis activation, leading to elevated glucocorticoid levels that can cross the placenta (14). After birth, caregiver mental health shapes the capacity to provide responsive care (48). Infant screen exposure is an increasingly common environmental input for the developing brain in this highly experience-dependent period (49).

### Social Exposures

Social exposures have been mostly studied through socioeconomic status (SES), a multidimensional construct spanning income, education, and occupation (50), offering several plausible pathways. Prenatally, it may act through associated factors like maternal nutrition and stress, and postnatally through parenting, access to resources, environmental health, neighborhood, and schooling characteristics (51).

## Methods

### Search strategy

This review was conducted in accordance with the Preferred Reporting Items for Systematic Reviews and Meta-Analysis (PRISMA) guidelines (52). We conducted a systematic search for eligible articles on November 17, 2025, in two databases: PubMed and Web of Science. The main components of the search were terms reflecting non-genetic exposures in the prenatal and early postnatal period, brain development assessed via magnetic resonance imaging (MRI), and neurodevelopmental outcomes. Full search strategies for all databases are available in the Supplementary Information. The reference lists of the included studies were manually screened to identify relevant records not captured in the electronic search. Two studies published after the search date were identified during writing. The review protocol was preregistered on the Open Science Framework (OSF; https://osf.io/q54rm).

### Selection criteria

#### Inclusion criteria

Studies were eligible if they (1) were original, peer-reviewed empirical studies published in English between January 2000 and March 2026; (2) were conducted in humans; (3) examined at least one non-genetic (biological, psychological, or social) exposure between conception and 3 years of age; (4) included at least one measure of brain morphology (structural MRI (sMRI) or diffusion MRI (dMRI)) acquired at any timepoint; and (5) reported at least one neurodevelopmental outcome assessed at any timepoint.

### Exclusion criteria

Studies were excluded if they were not original empirical reports, relied solely on retrospective parental recall of the exposure collected more than one year after it without corroborating records, were intervention studies, focused on genetic diagnoses or neurodevelopmental disorders as the primary influence, used a categorical operationalization without a control group, lacked sufficient methodological detail, or did not correct for multiple comparisons.

### Study identification

Study identification began by removing duplicate records using Rayyan (53). Two authors (JD and HB) independently reviewed the titles and abstracts of the papers prior to full-text screening. Disagreements were resolved by a third author (RB). One author (JD) then assessed full texts for eligibility.

### Data extraction and risk of bias assessment

Relevant information for each study included sample characteristics; the exposure and its timing; MRI modality, timing, measures, and brain regions; and neurodevelopmental outcome concepts and timing. We further extracted the associations among exposure, brain, and outcome – including mediation and sex-difference analyses when conducted. If reported and applicable, we extracted secondary datasets, country, birth cohort, and parental demographics. All studies underwent risk of bias assessments using the Risk of Bias in Non-Randomized Studies of Exposure (ROBINS-E) tool (54).

### Data synthesis and visualization

Studies investigating the same concept were grouped into subcategories. Concepts examined by two or more studies were integrated into the results, discussion, and visualizations; concepts examined by only a single study or as a non-decomposable composite are not further discussed but described in Table S1 (55–59).

To summarize structural findings and their consistency across studies, results from the included studies were visualized on brain templates using a structured scoring approach. Briefly, for each exposure, indicators of. delayed vs. accelerated brain maturation were counted region-wise (e.g., lower vs. higher volumes, cortical thickness, or white matter integrity; for full mapping see Table 1). Cortical regions were defined at the lobe-level; subcortical structures and white matter tracts were abstracted from the included studies (see Figure 2). For each combination of exposure and brain region, counts for delayed vs. accelerated maturation were summarized with a Laplace-smoothed score bounded between −1 (consistent delayed maturation) and +1 (consistent accelerated maturation). Additionally, a frequency count of how often these structural changes of accelerated or delayed maturation were associated with favorable or unfavorable neurodevelopmental outcomes was established. Further details on score derivation, template generation, and visualization are provided in the Supplementary Information and Tables S4–S10.

**Table 1:**
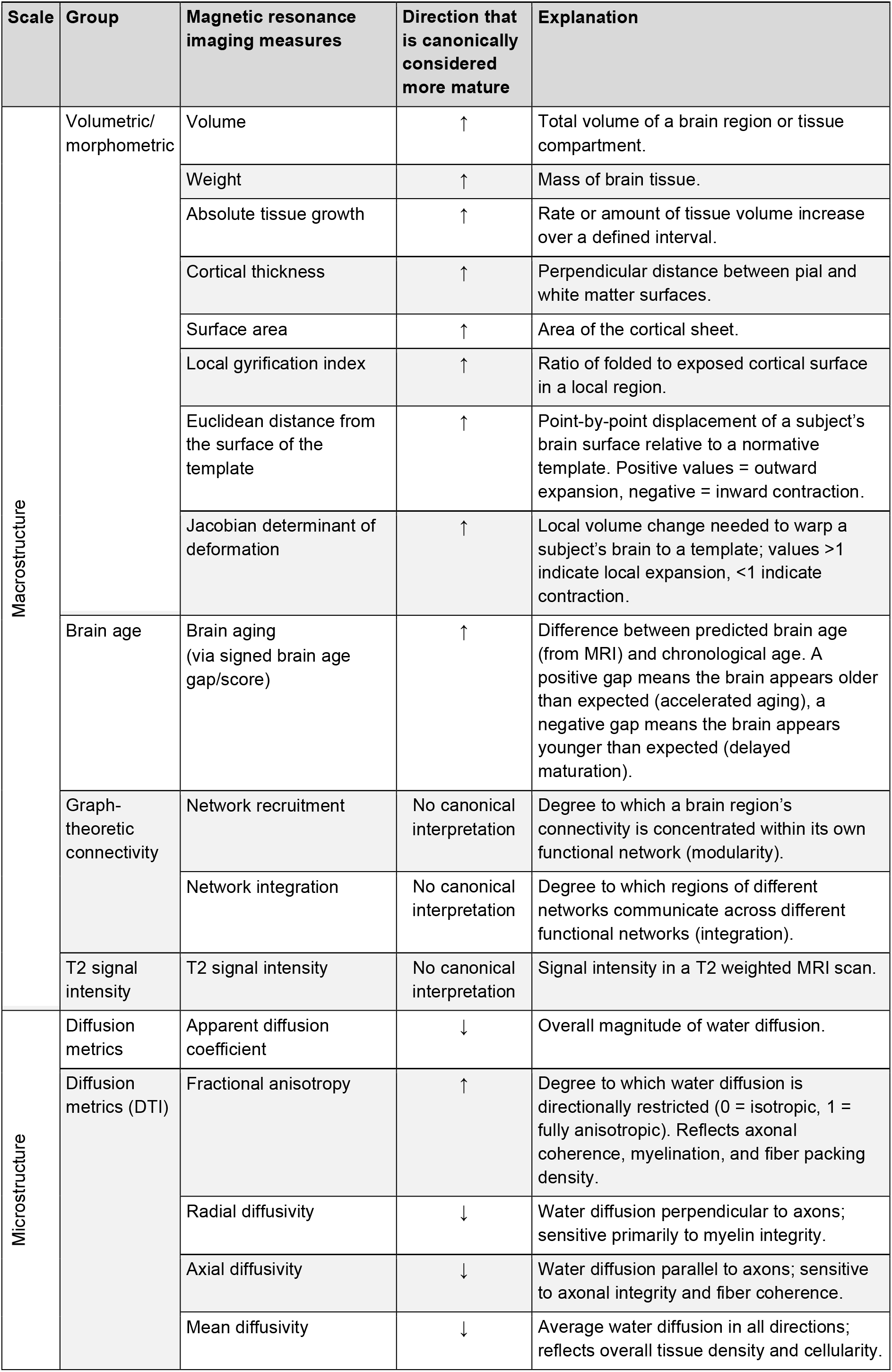
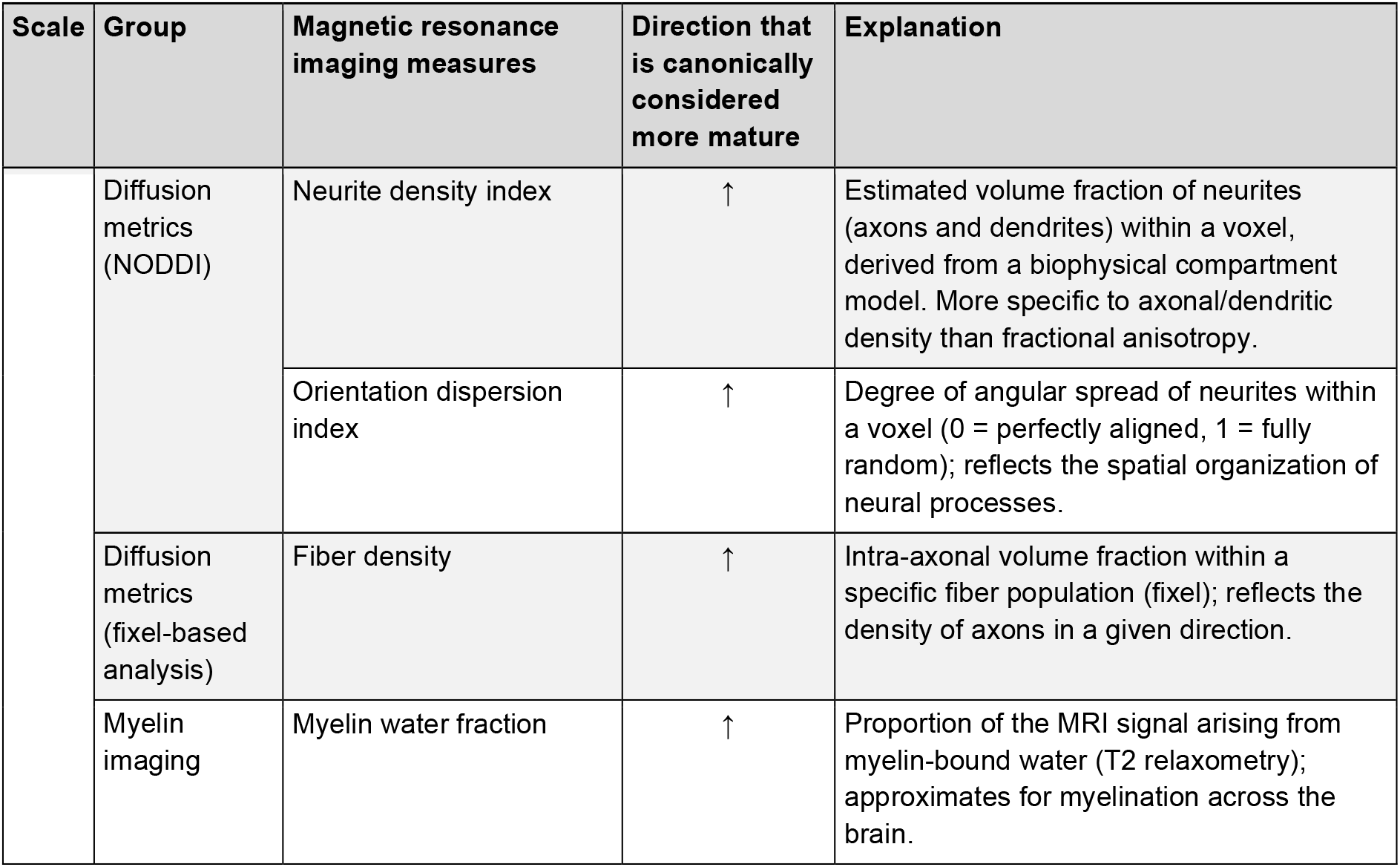
MRI measures used in the studies included. Extrapolating over- or under-maturation from an imaging phenotype is not always straightforward; directional interpretations and brief explanations were therefore based on the terminology used in the included studies and are applied consistently throughout this review. For microstructural measures, higher maturation can generally be interpreted as higher microstructural integrity or organization. DTI, diffusion tensor imaging; MRI, magnetic resonance imaging; NODDI, neurite orientation dispersion and density imaging.

**Figure 2:**
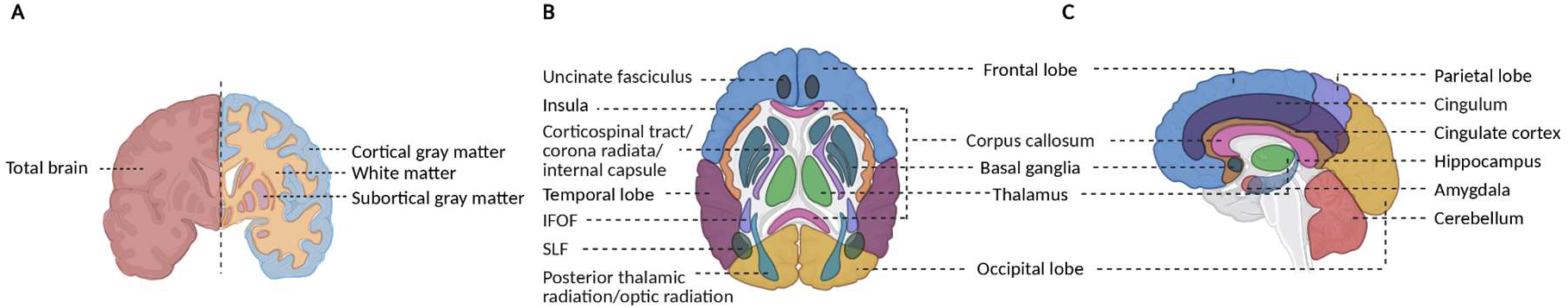
Template for Laplace-score visualization with (**A**) global measures and (**B**, **C**) regional measures. IFOF, inferior fronto-occipital fasciculus; SLF, superior longitudinal fasciculus I, II, and III. This figure was created in BioRender, https://BioRender.com.

## Results

### Study selection

Our initial search yielded 507 records (PubMed: 269; Web of Science: 238). After removing duplicates, 448 records underwent title and abstract screening; 391 were excluded, leaving 57 for full-text screening. Thereof, 40 articles met eligibility criteria. Additionally, reference-list screening and ongoing monitoring of the literature revealed 15 more eligible articles (Table 2; Figure 3).

**Table 2:**
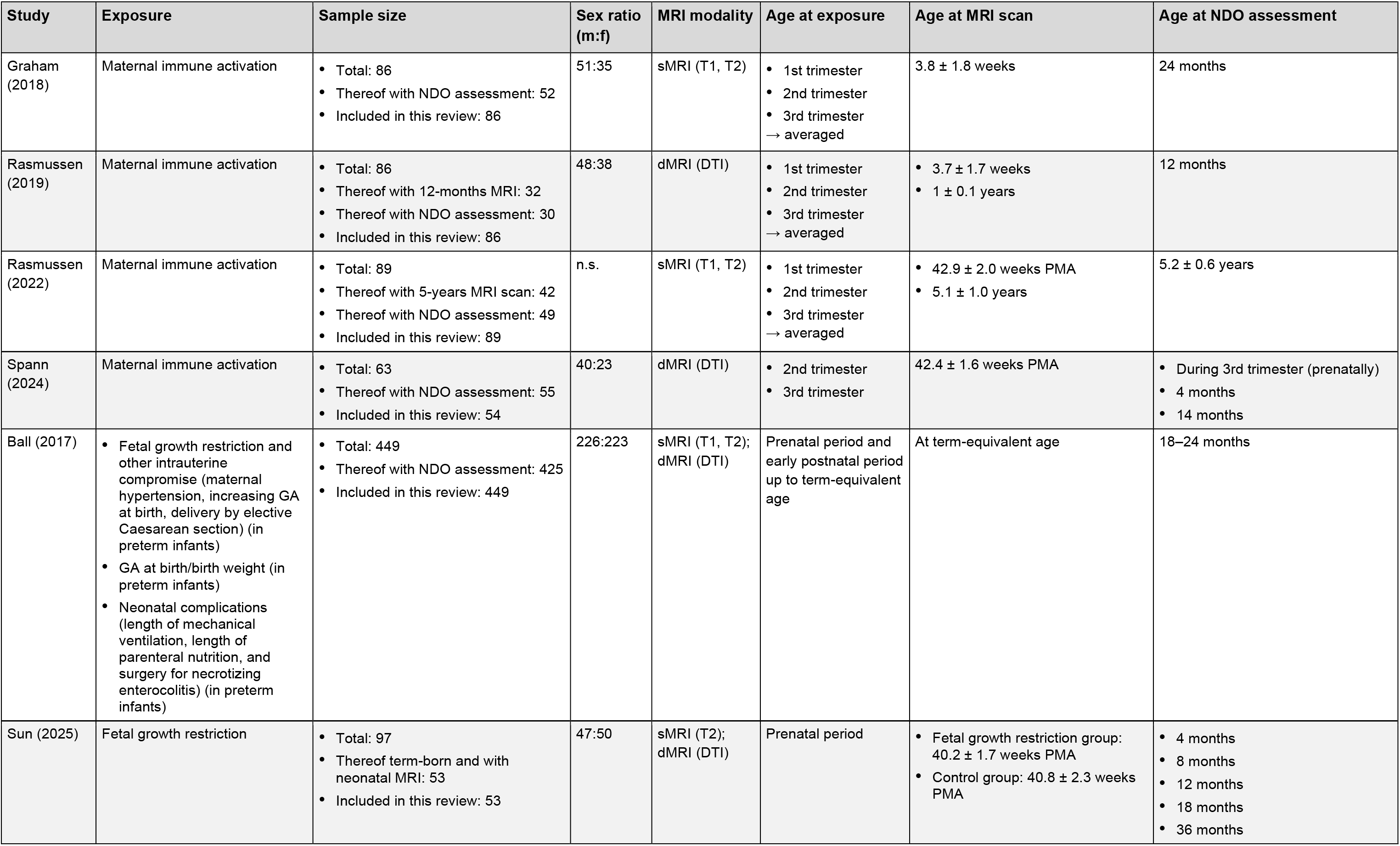
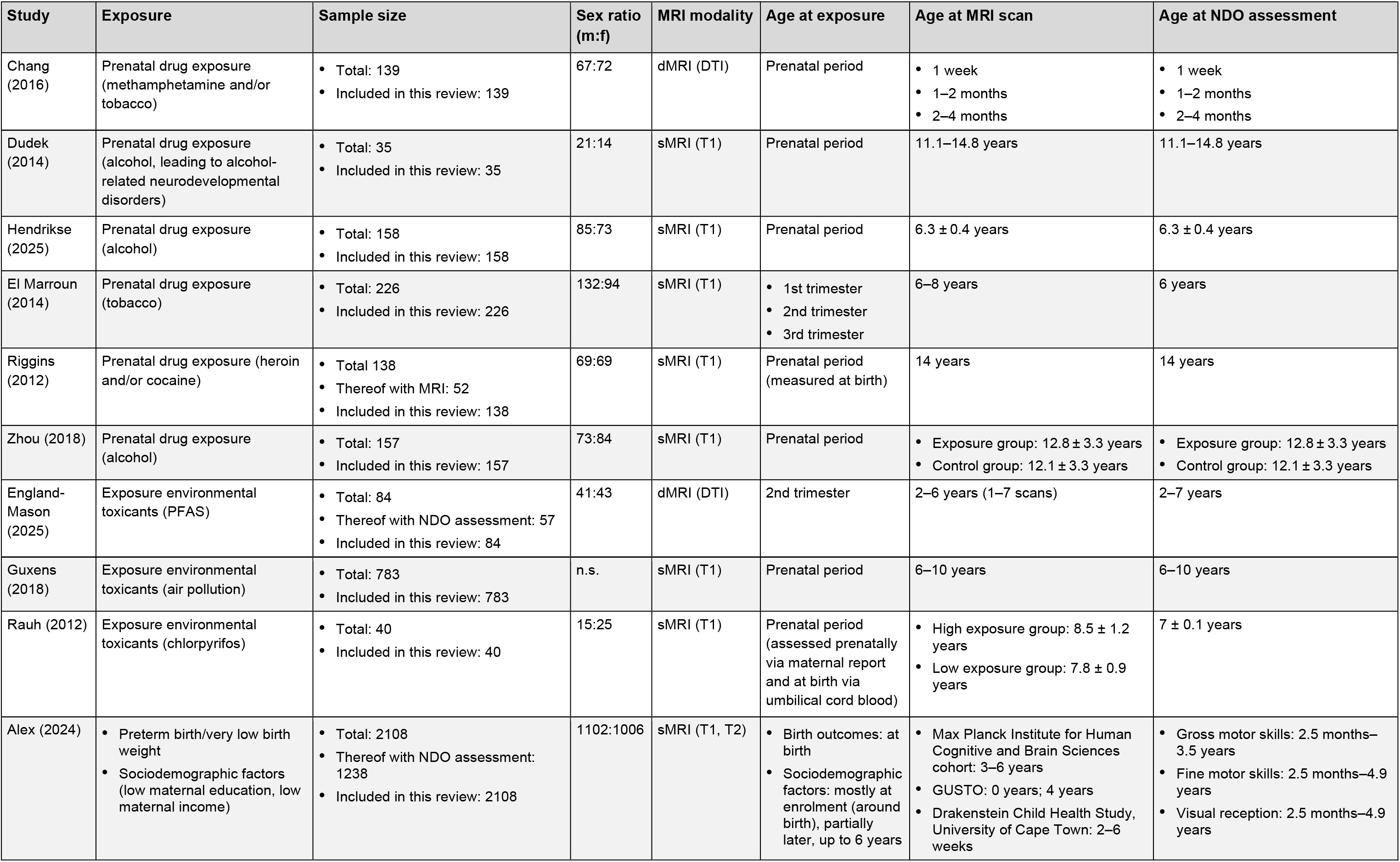
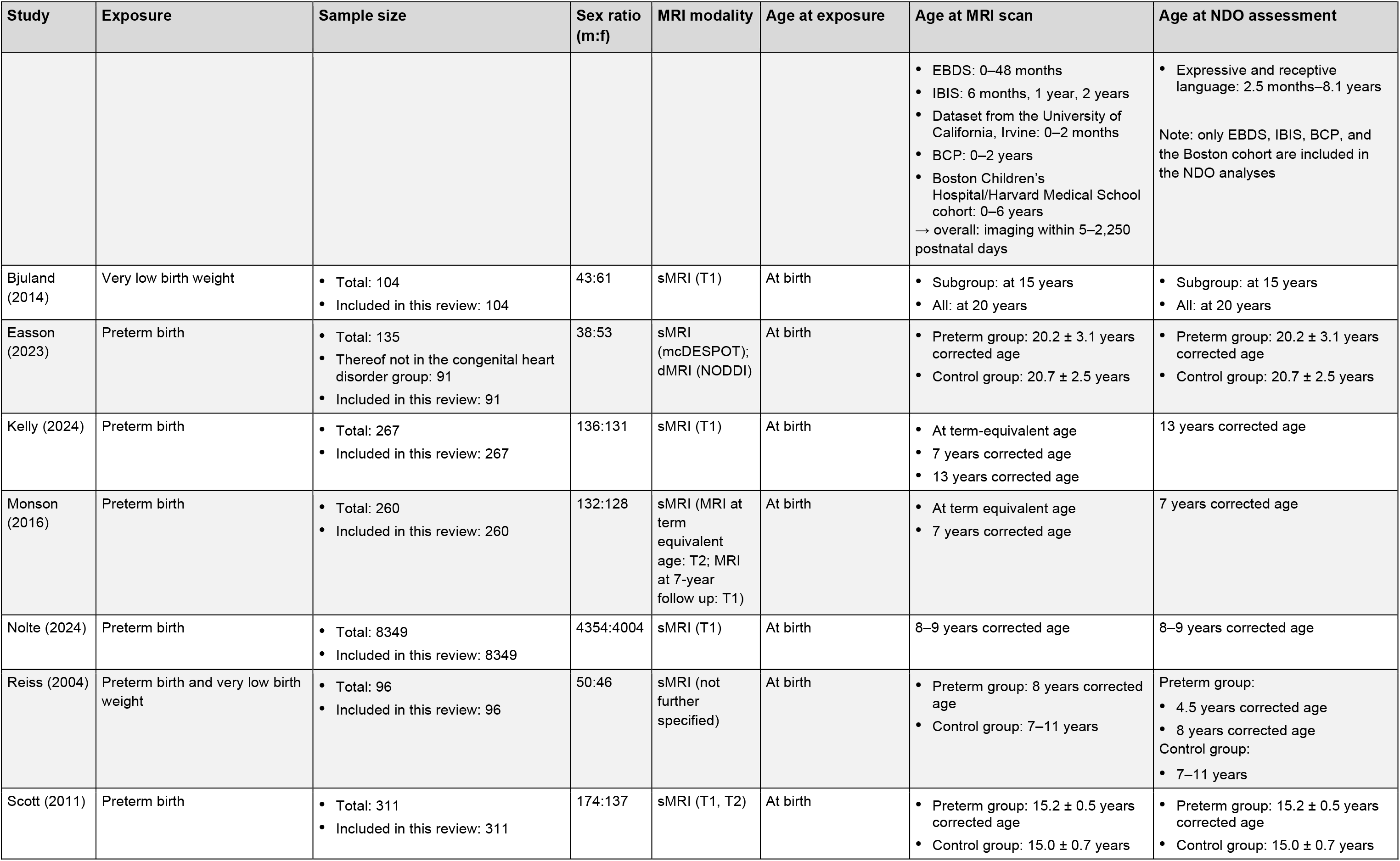
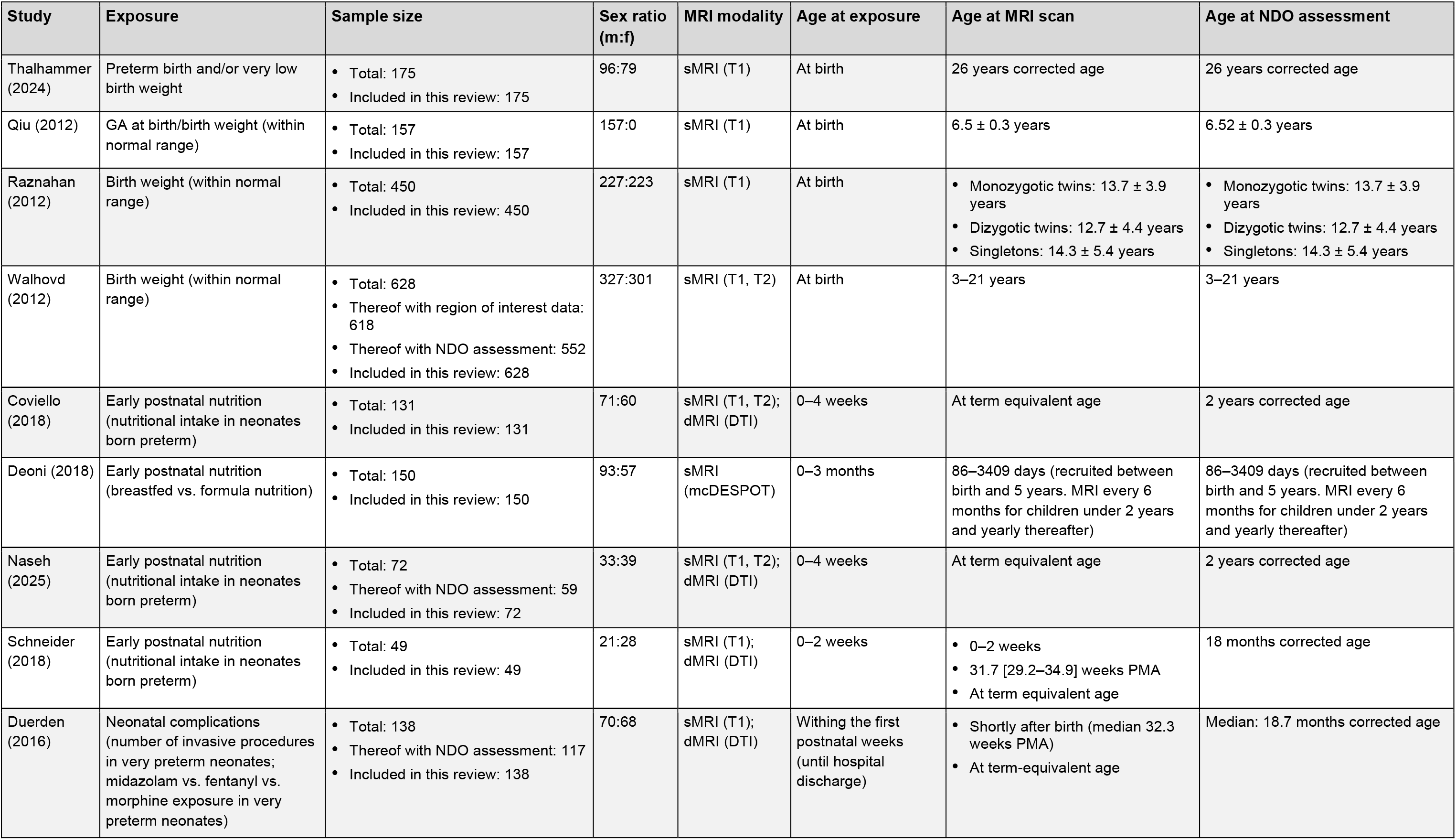
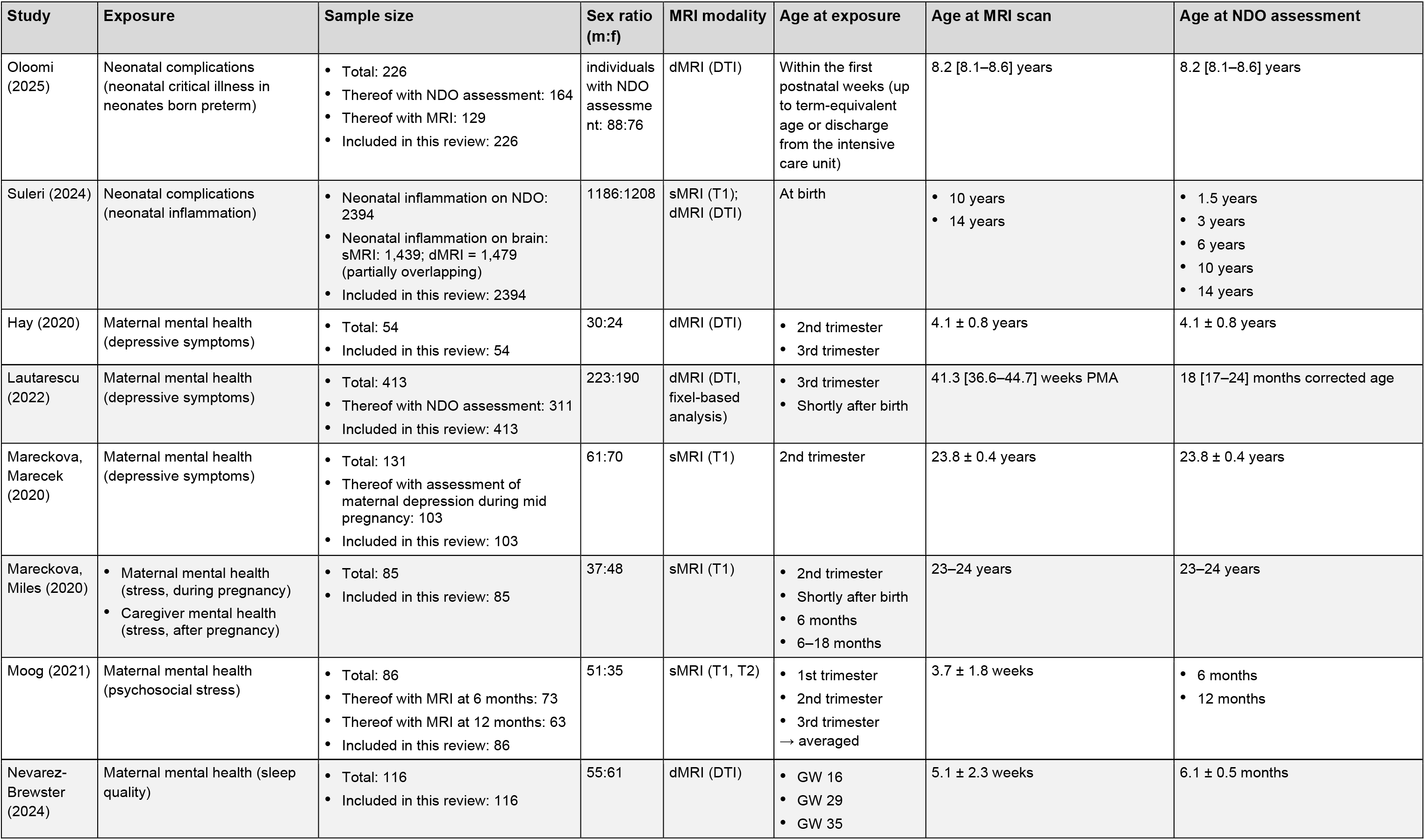
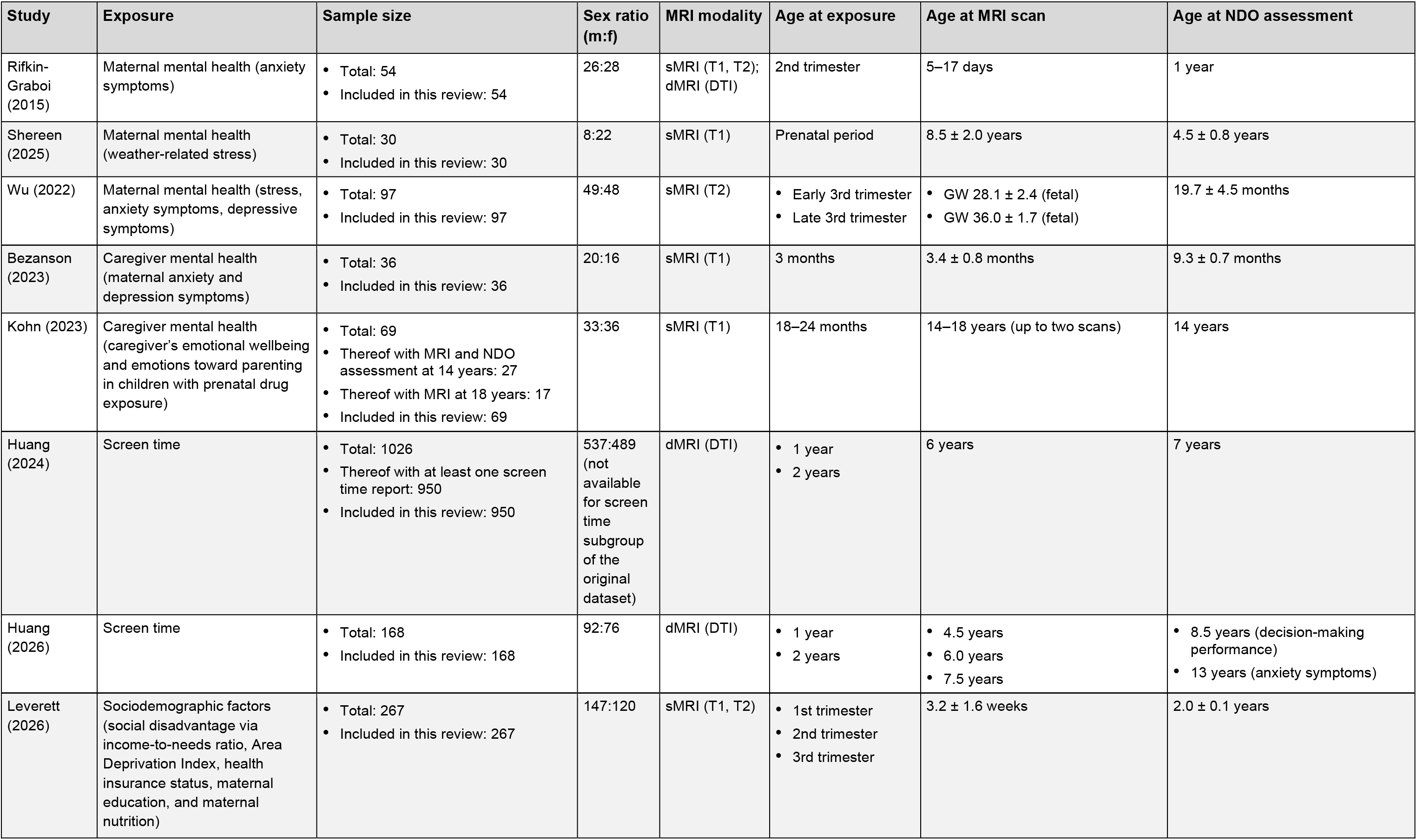
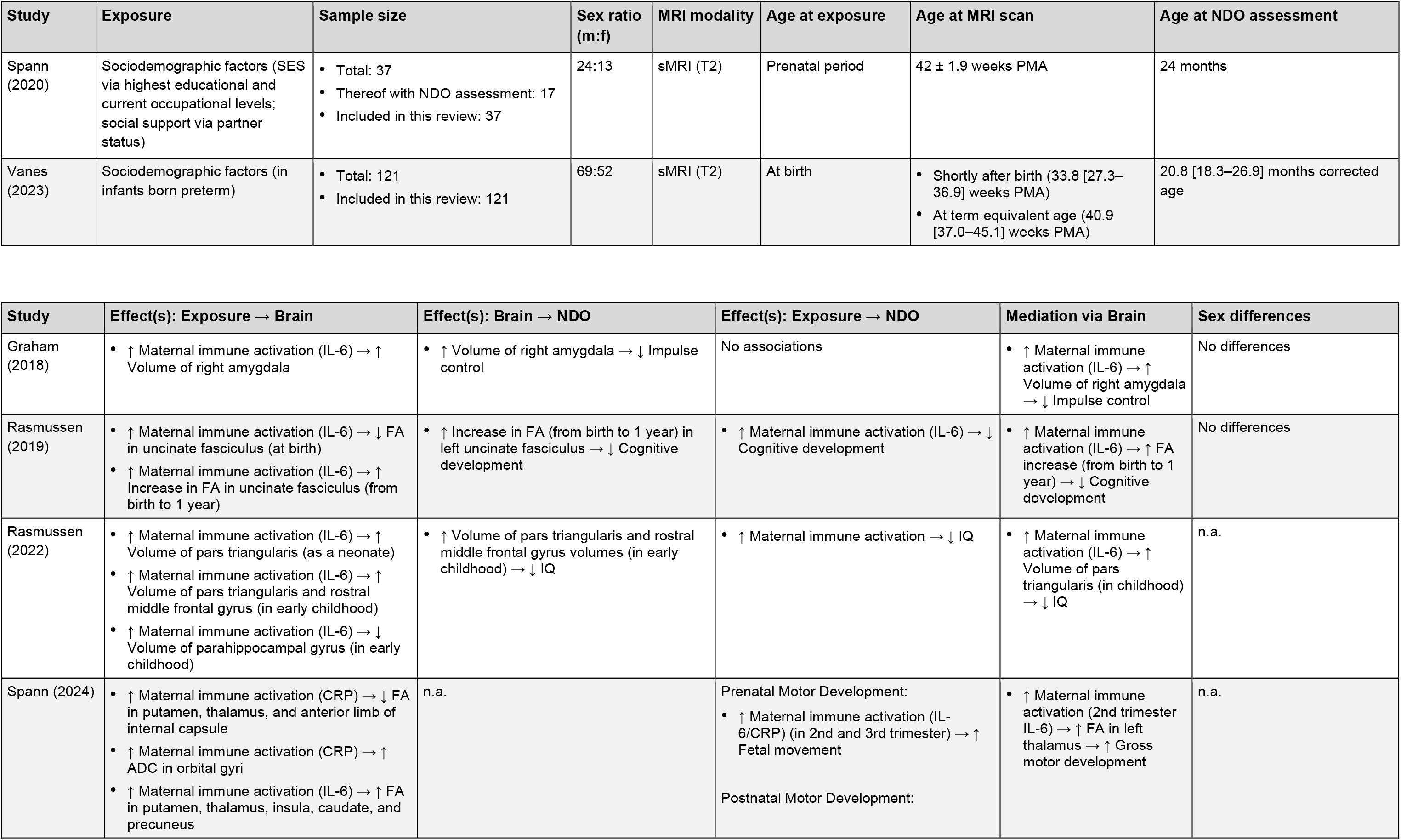
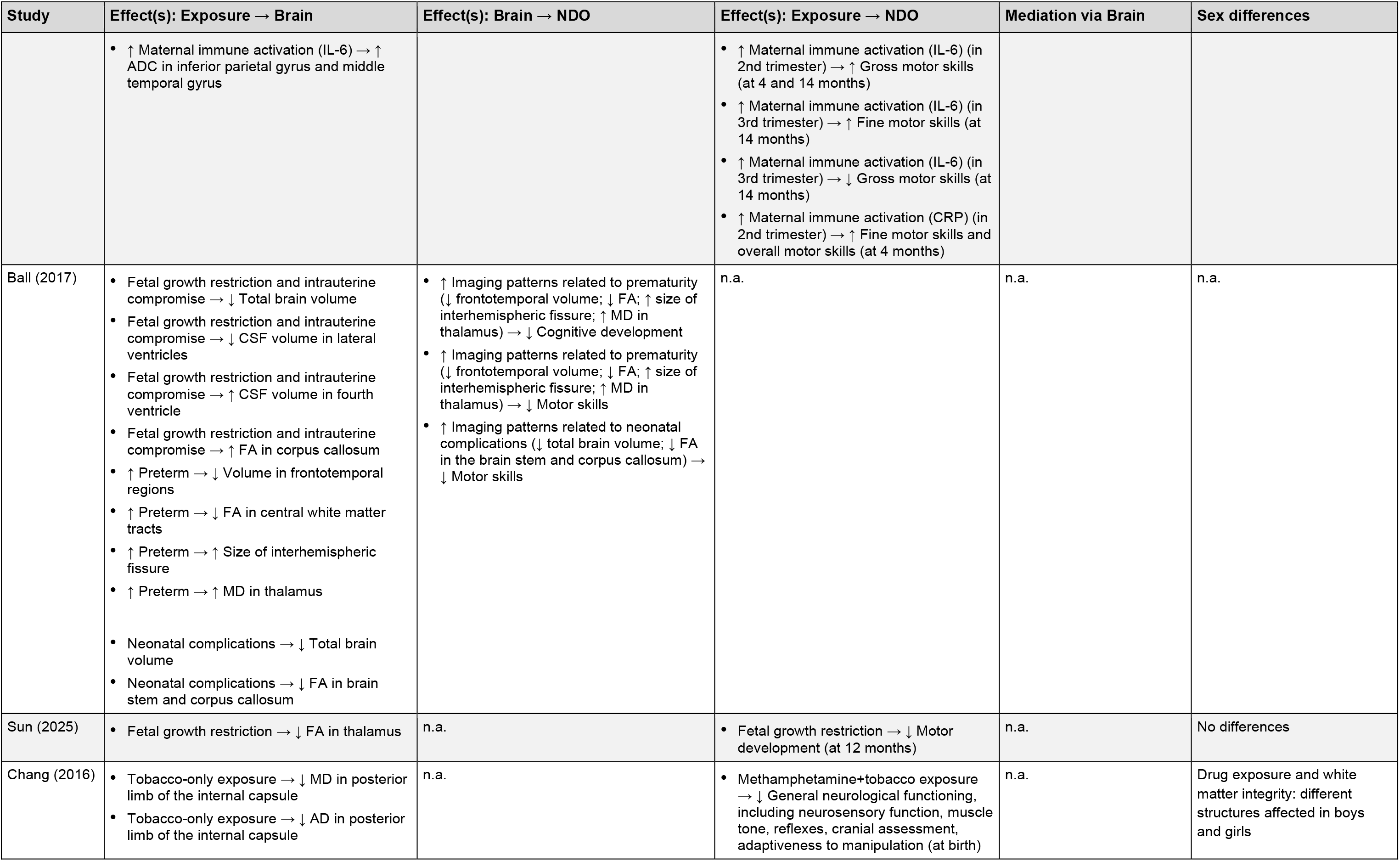
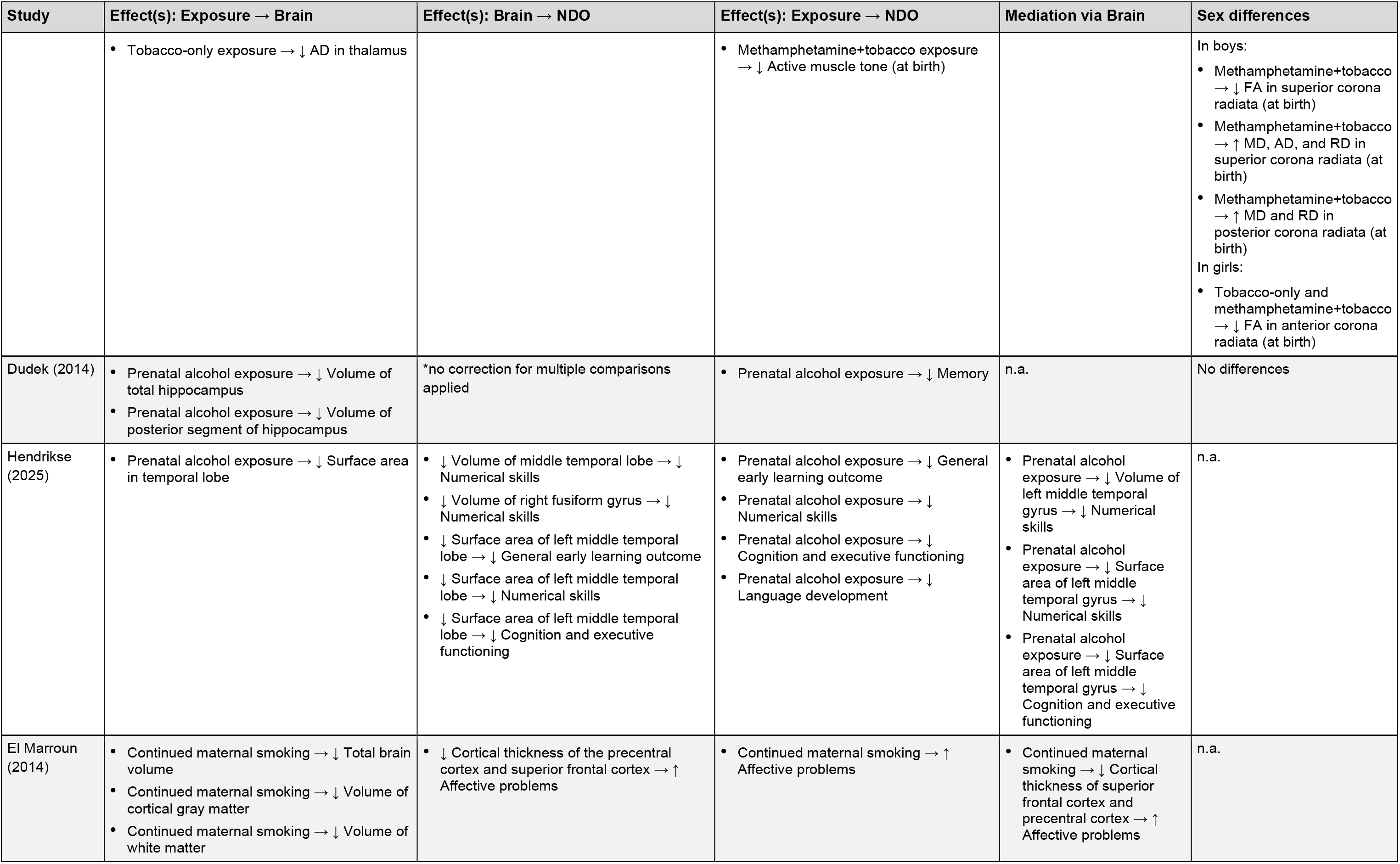
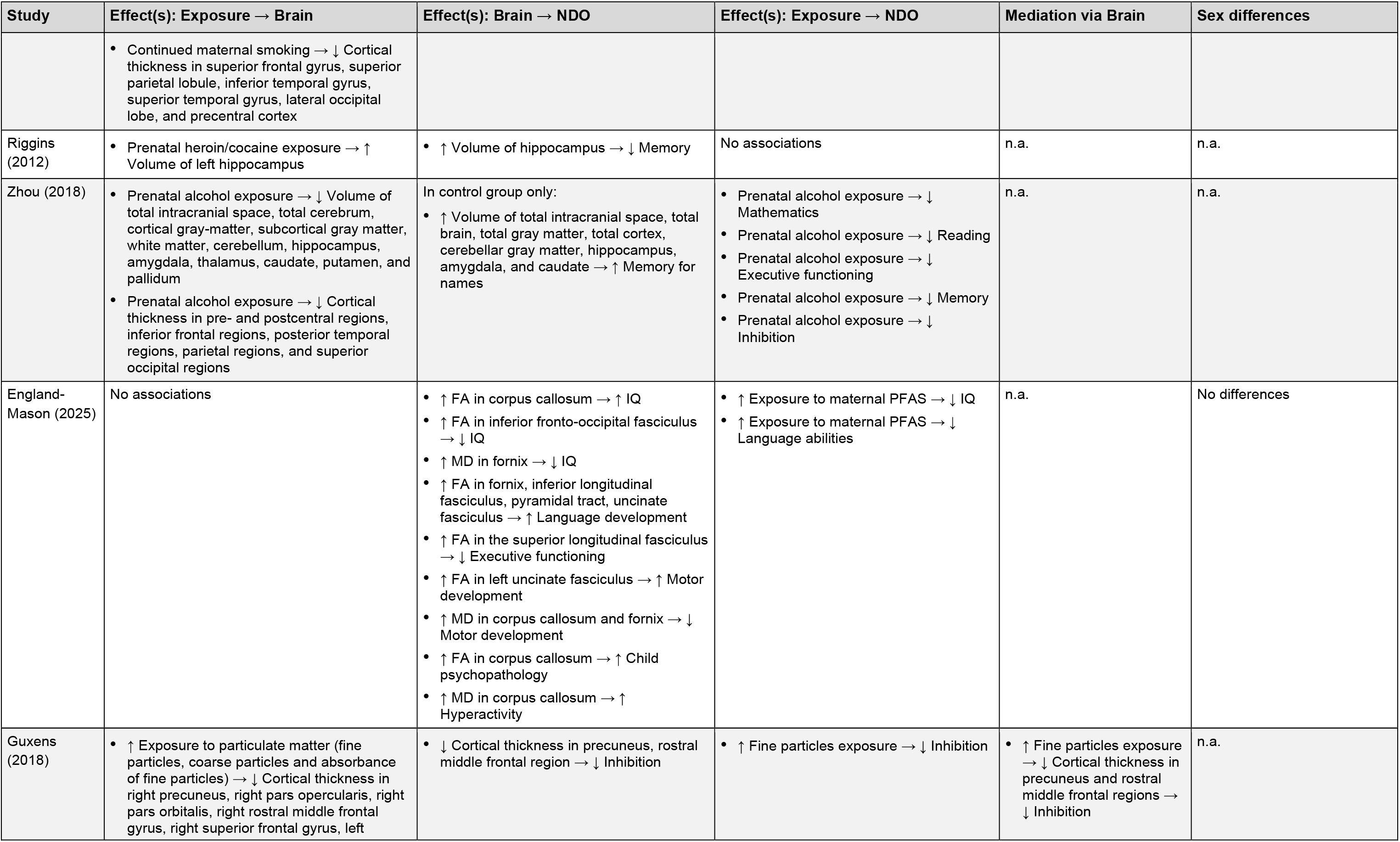
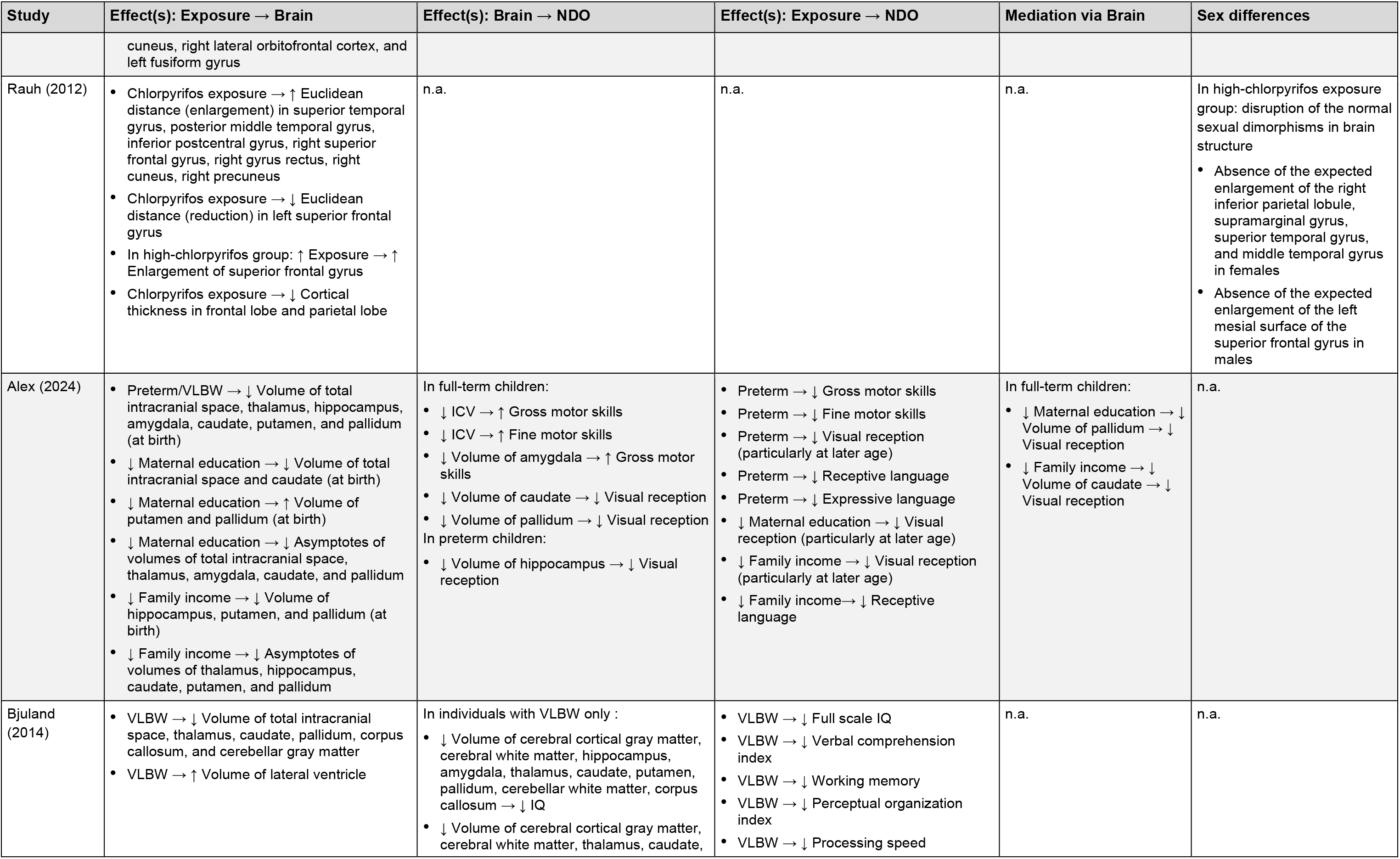
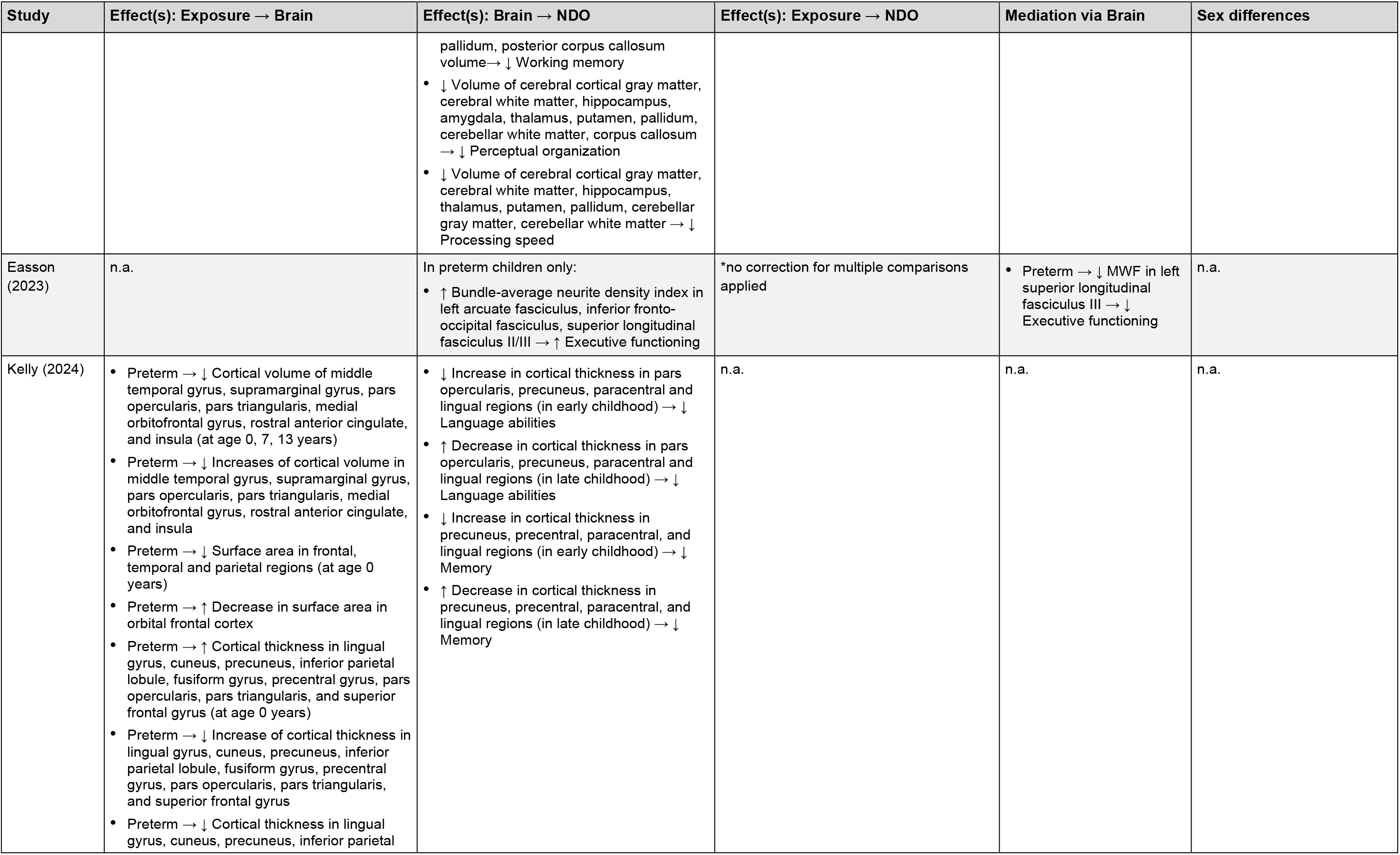
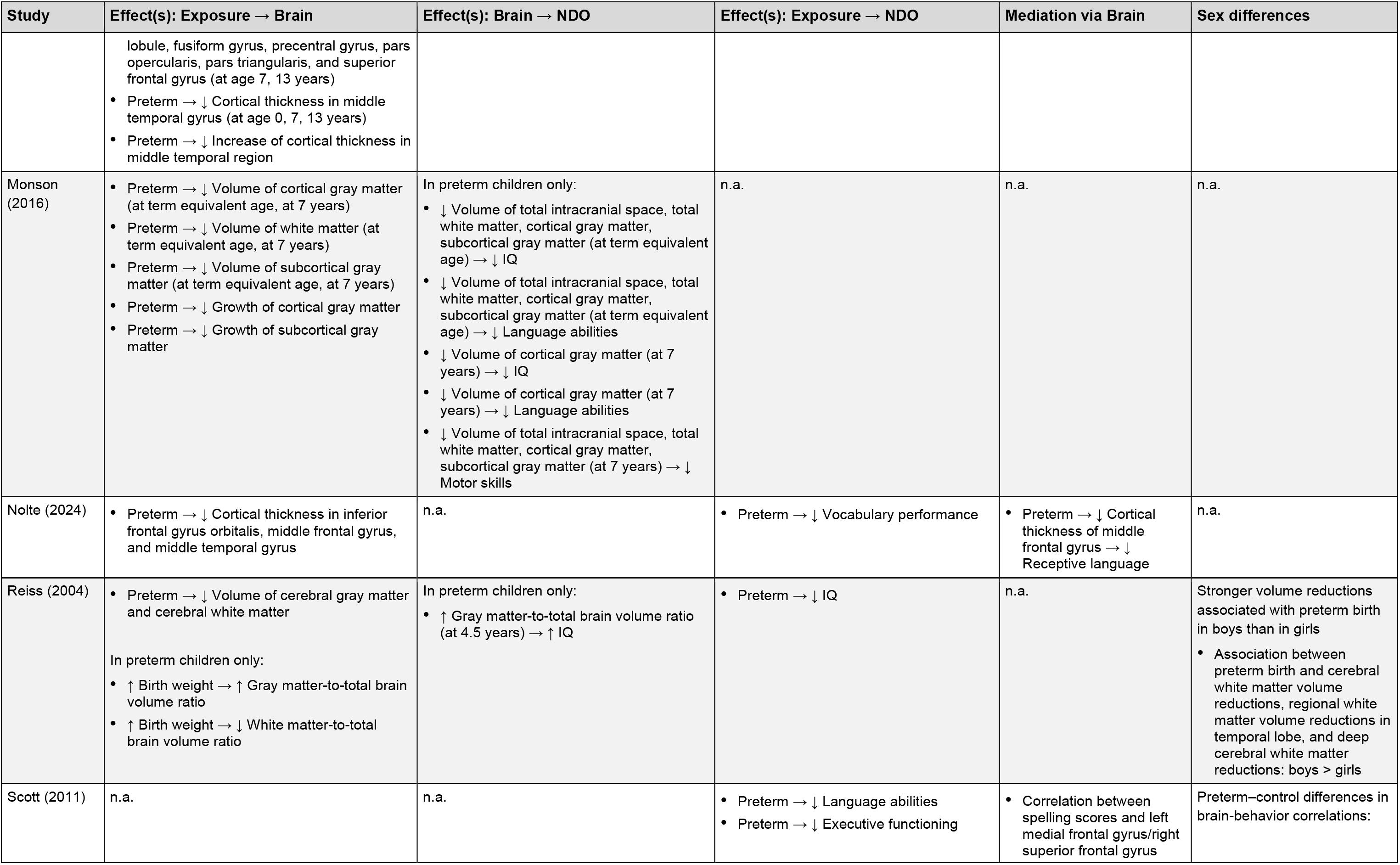
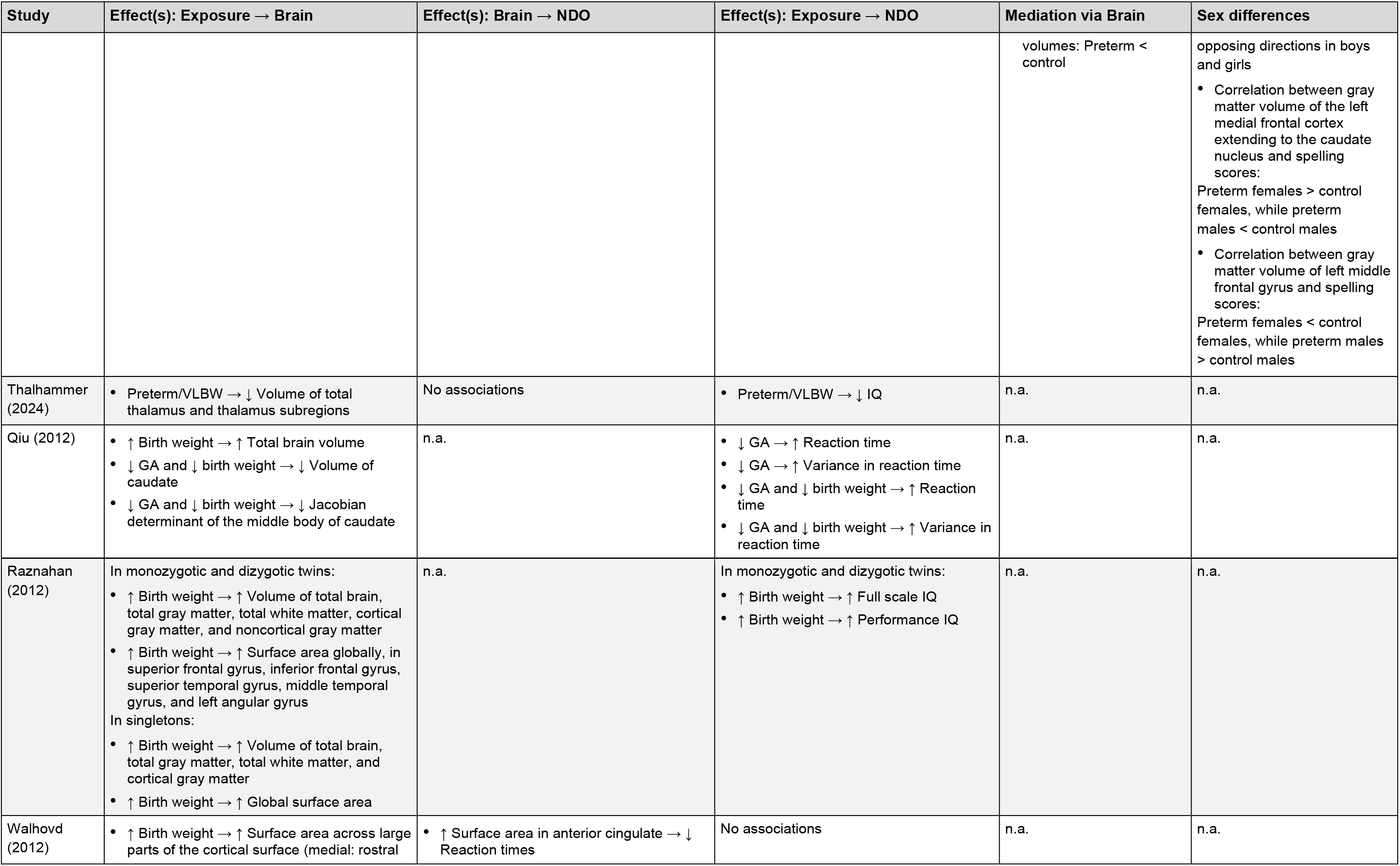
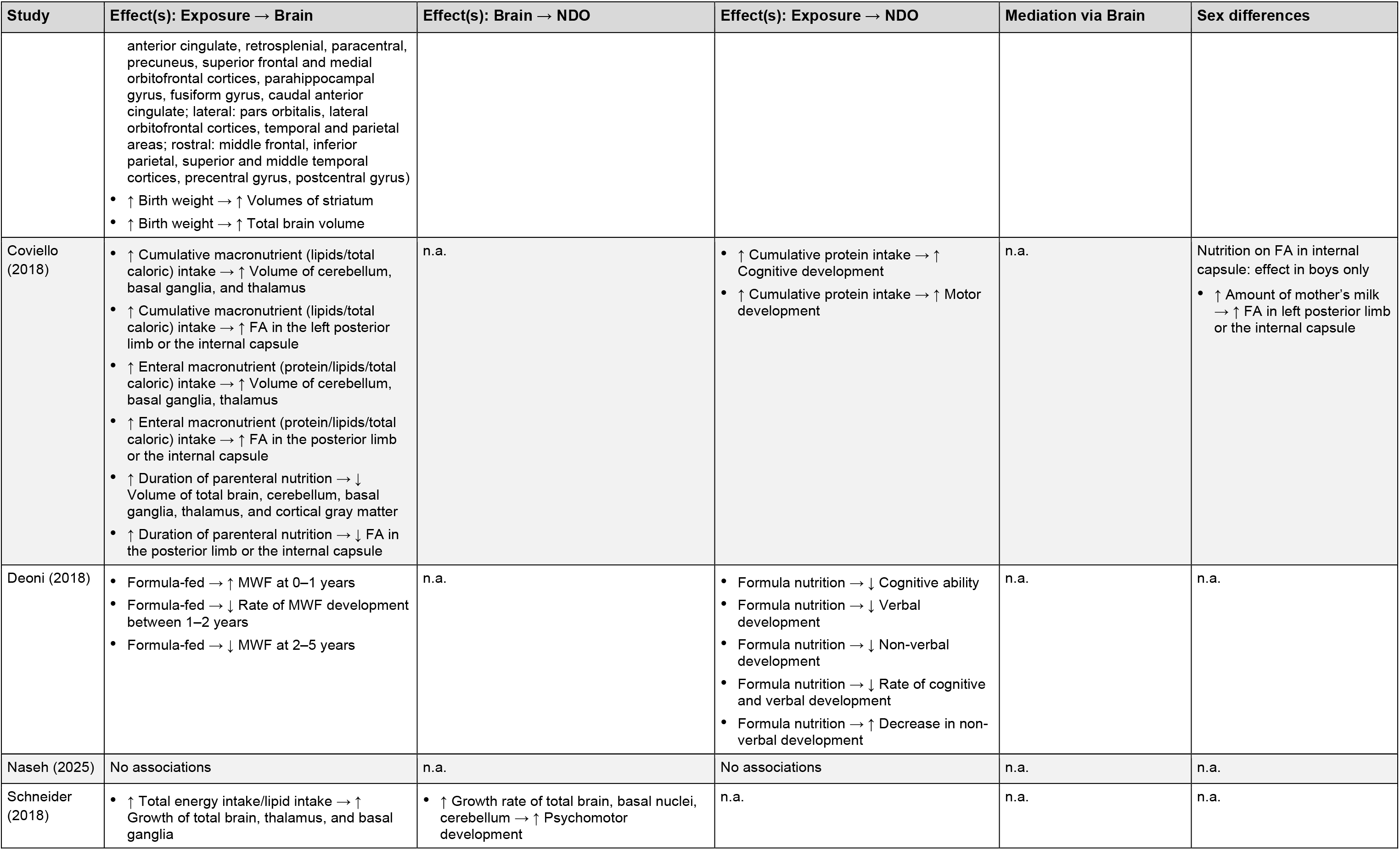
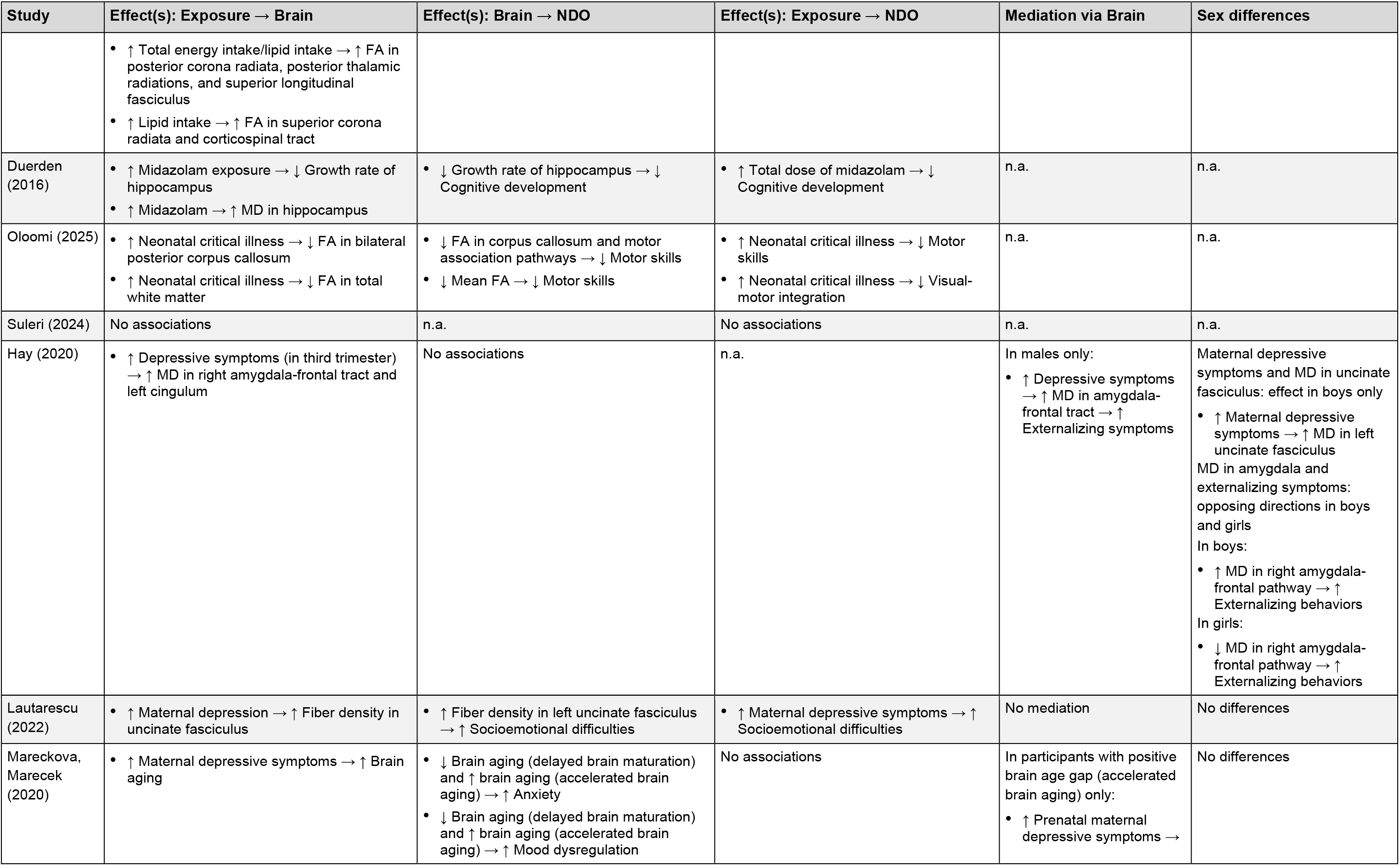
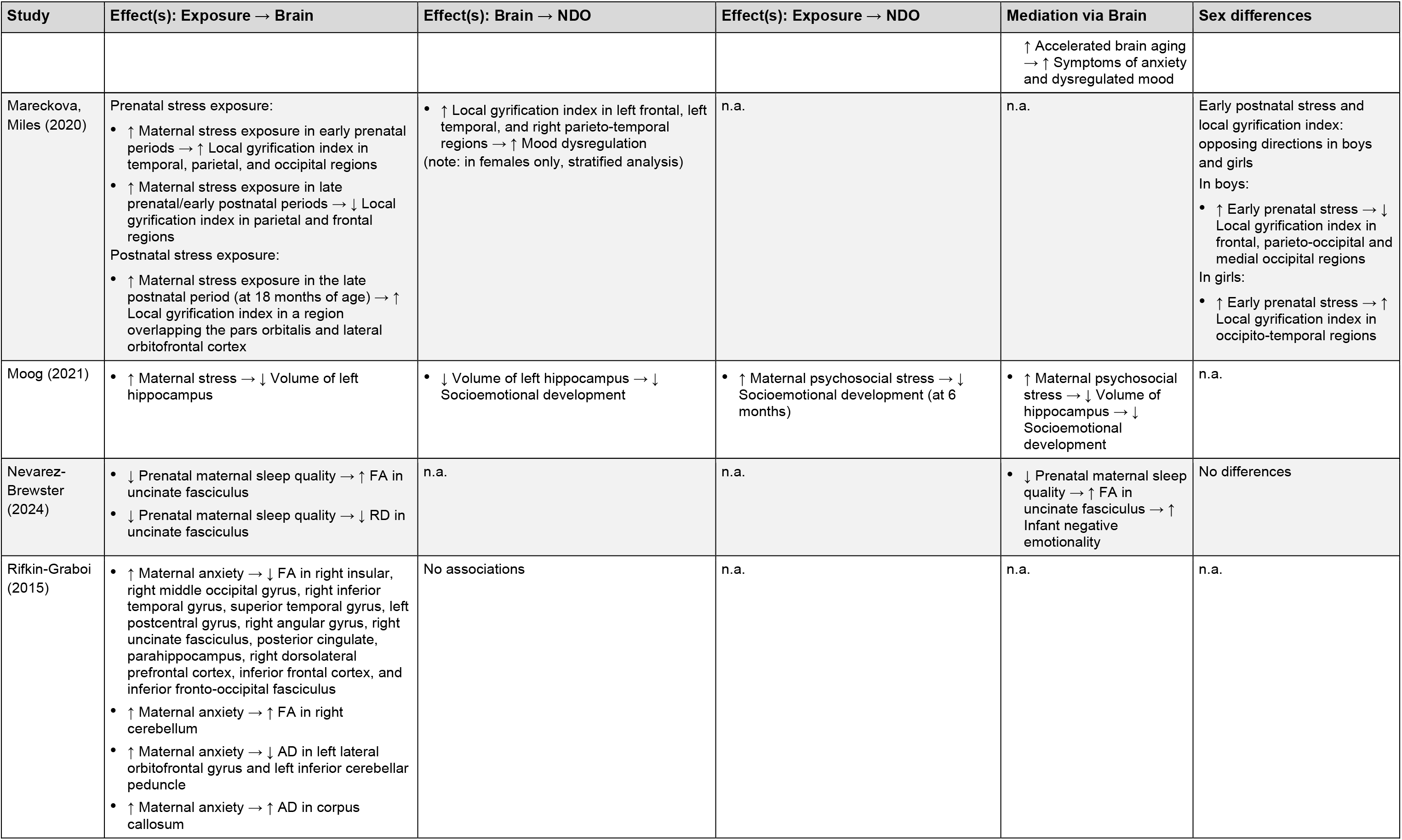
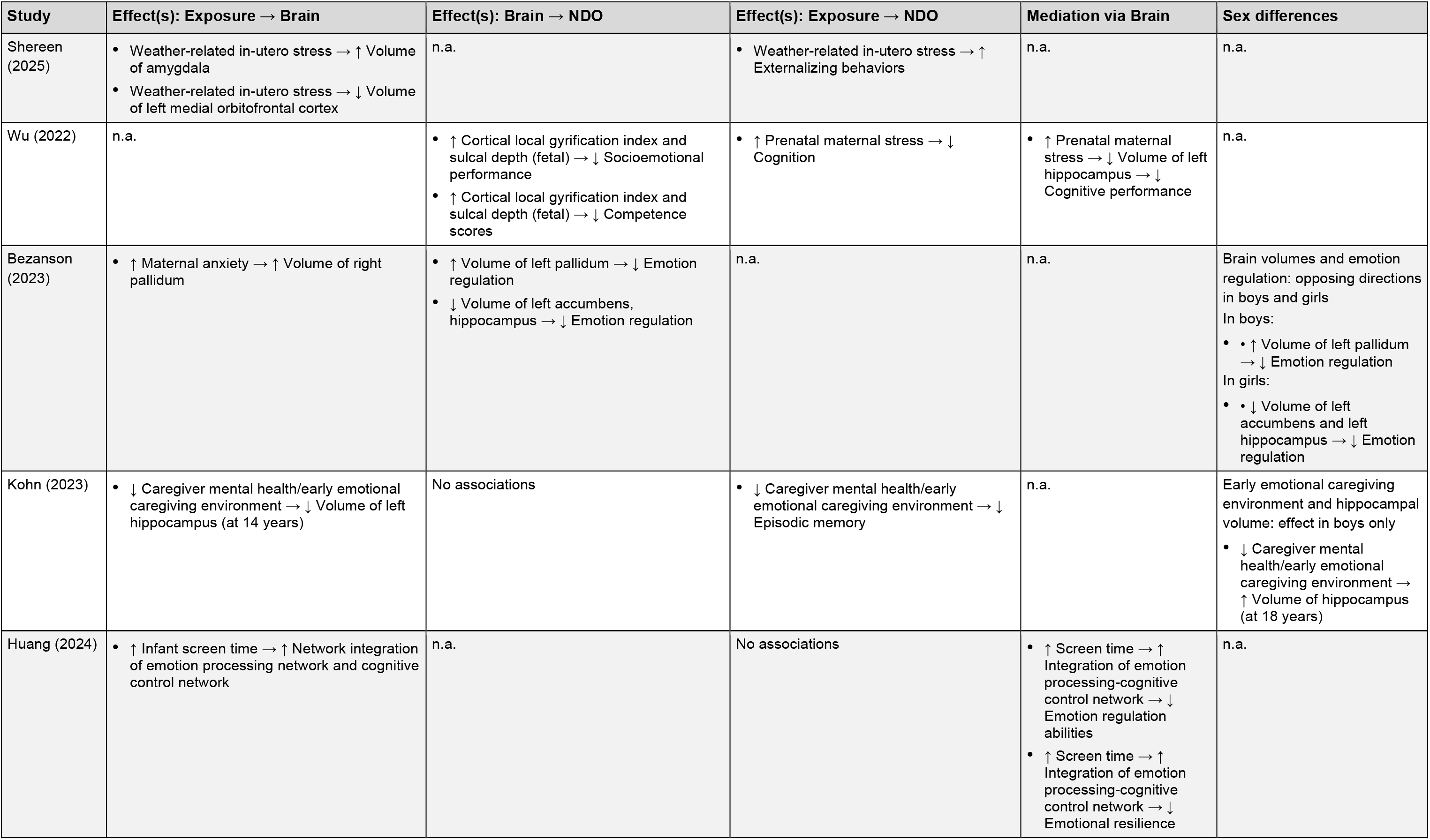
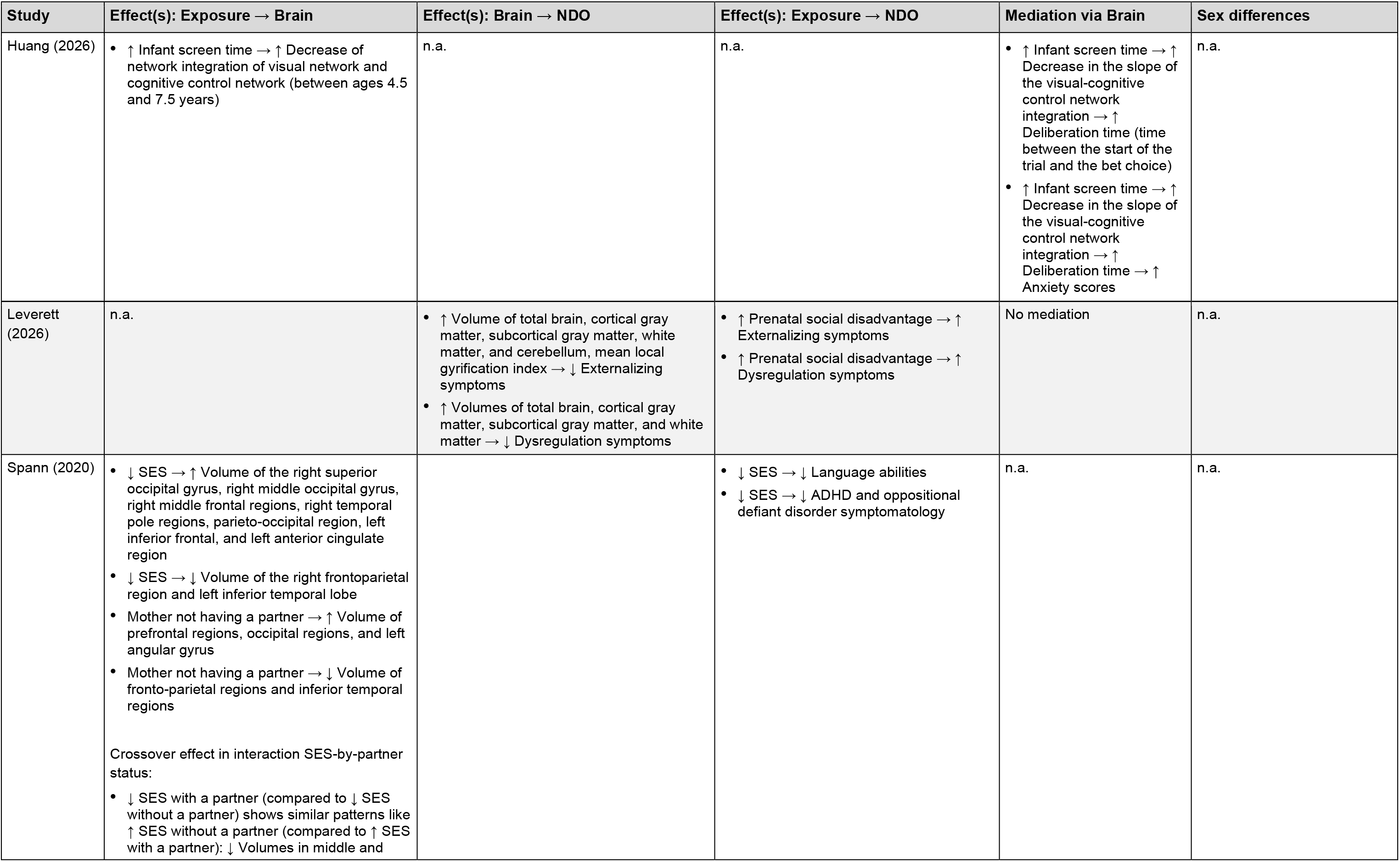
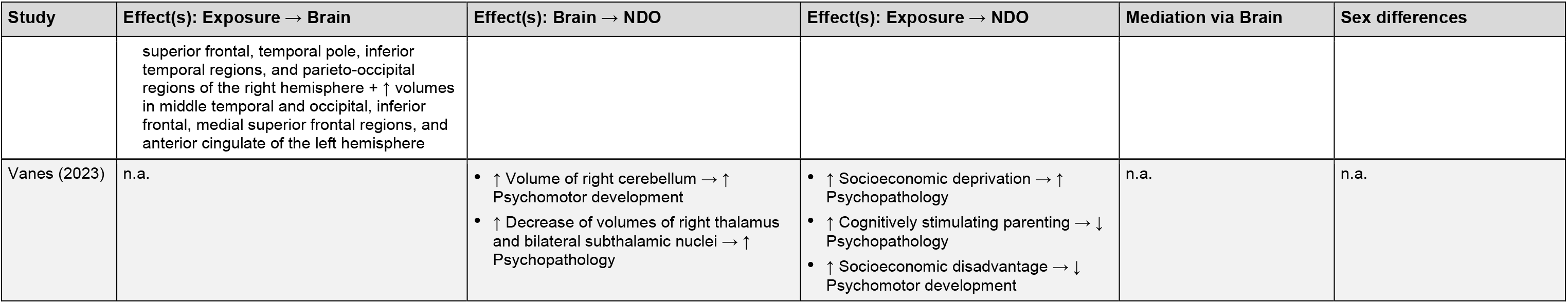
Overview of studies assessing the neurodevelopmental consequences of biological, psychological, and social early-life exposures. Results that did not survive correction for multiple comparisons or became non-significant after covariate inclusion are treated as non-significant. Age information represents postnatal age, if not stated otherwise, in mean ± SD, mean [range], or range (depending on the available information), if not stated otherwise. Further information on the studies can be found in Table S2. AD, axial diffusivity; ADC, apparent diffusion coefficient; ADHD, attention deficit hyperactivity disorder; BCP, Baby Connectome Project; CRP, C-reactive protein; CSF, cerebrospinal fluid; dMRI, diffusion MRI; DTI, diffusion tensor imaging; EBDS, Early Brain Development Study; FA, fractional anisotropy; GA, gestational age; GUSTO, Growing Up in Singapore Towards Healthy Outcomes; GW, gestational week(s); IBIS, Infant Brain Imaging Study; ICV, intracranial volume; IL-6, interleukin-6; IQ, intelligence quotient; mcDESPOT, Multicomponent Driven Equilibrium Single Pulse Observation; MD, mean diffusivity; MRI, magnetic resonance imaging; MWF, myelin water fraction; n.a., not applicable; NDO, neurodevelopmental outcome(s); NODDI, neurite orientation dispersion and density imaging; n.s., not specified; PFAS, per- and polyfluoroalkyl substances; PMA, postmenstrual age; RD, radial diffusivity; SES, socioeconomic status; sMRI, structural MRI; VLBW, very low birth weight.

**Figure 3:**
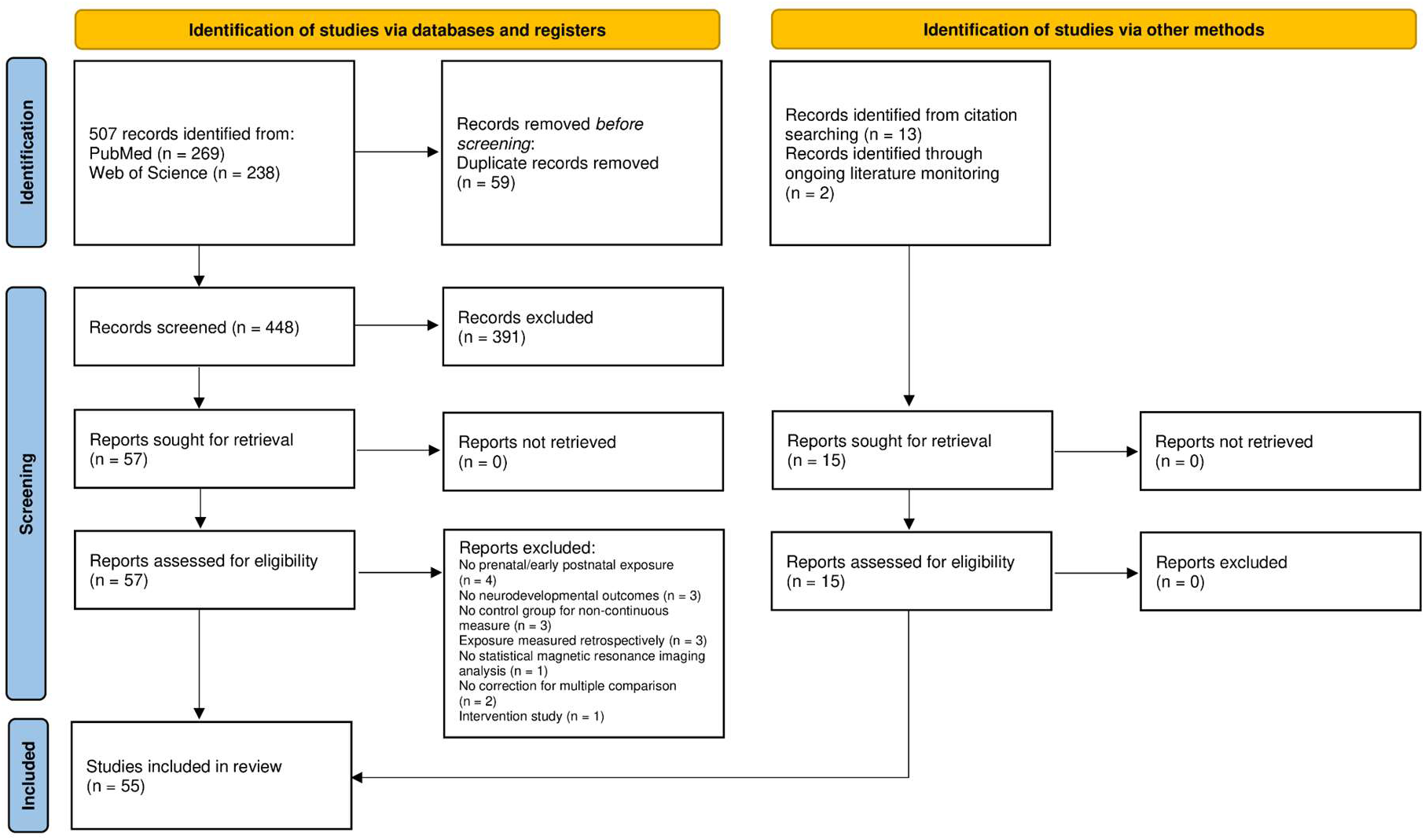
PRISMA 2020 flow diagram of the study selection process.

### Study characteristics

The included studies span from 2004 to 2026, with only one study published prior to 2011. Studies after 2020 were more likely to focus on psychological and social exposures (see Figure 4).

**Figure 4:**
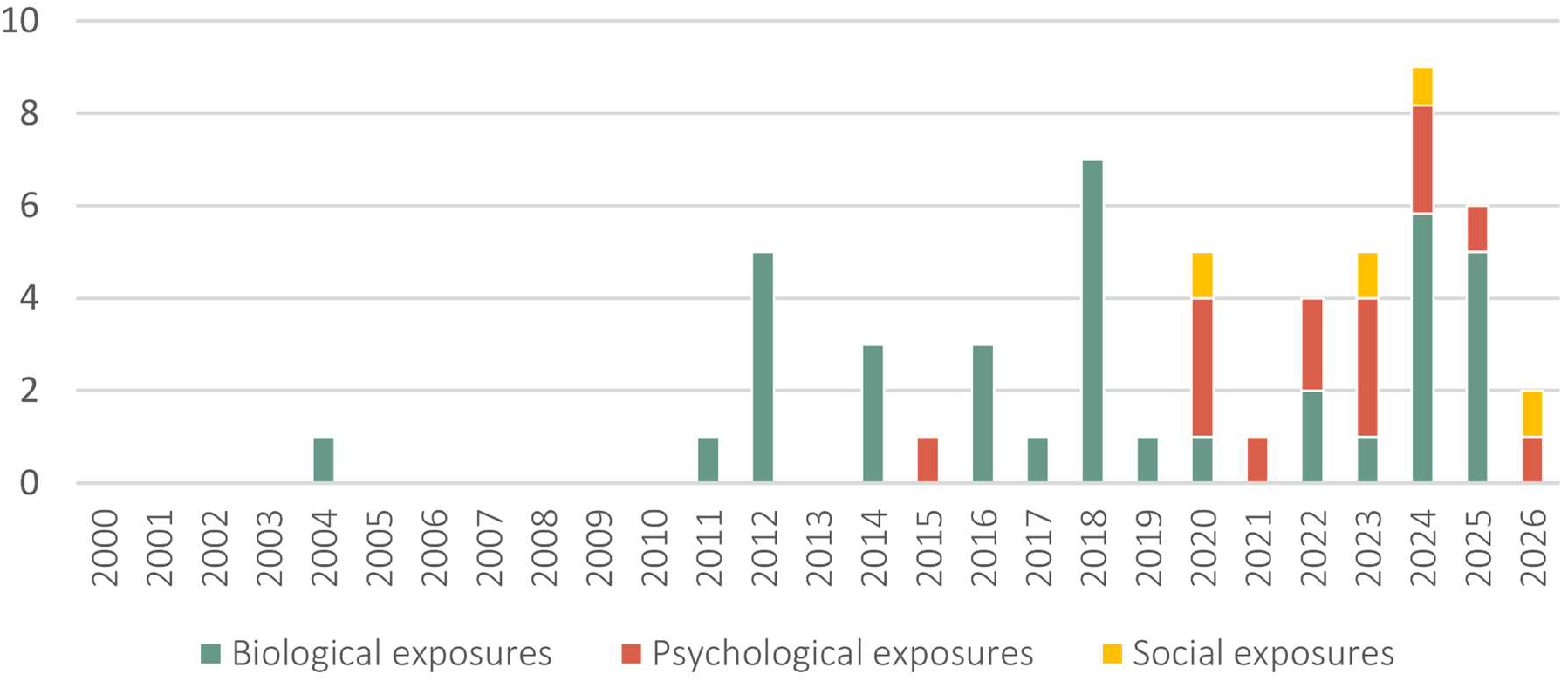
Publications included in this review over time. Number of papers per year, separated by exposure category (biological, psychological, and social exposures).

In total, 22,679 participants were included across studies, with a median sample size of n = 121; minimum n = 30 (60); maximum n = 2108 (61). Note that some participants may be counted more than once, as several studies used the same secondary datasets (for metadata on all included studies and further details, see Supplementary Information and Table S2). The majority of studies were rated as having a low risk of bias or “some concerns”, see Table S3 for the full assessment.

Exposures were assessed via biospecimens, medical records, and questionnaires; brain morphology via sMRI and dMRI (Table 1); and neurodevelopmental outcomes via questionnaires, performance tests, and behavioral observations; see Supplementary Information and Table S2.

Most studies focused on a single exposure; a few investigated multiple exposures (45,61,62) and were therefore considered in all respective categories. One study (55) used a composite measure of biological, psychological, and social exposures and could not be assigned to a single category.

**Key findings: How Early-Life Exposures Shape Brain Structure and Neurodevelopmental Outcomes**

#### Biological Exposures

##### Prenatal

*Maternal immune activation* (n = 4; Figure 5, **A**). Higher prenatal maternal immune activation was largely associated with more mature macrostructure, particularly with higher volumes in the amygdala (63) and frontal regions (64). Higher volumes mediated links with lower impulse control and lower fluid intelligence. Microstructural findings were more mixed across regions, partly depending on the specific immune marker (65,66). Changes in both directions were mostly associated with poorer outcomes across cognition and executive functioning, with some beneficial motor effects (66).

**Figure 5:**
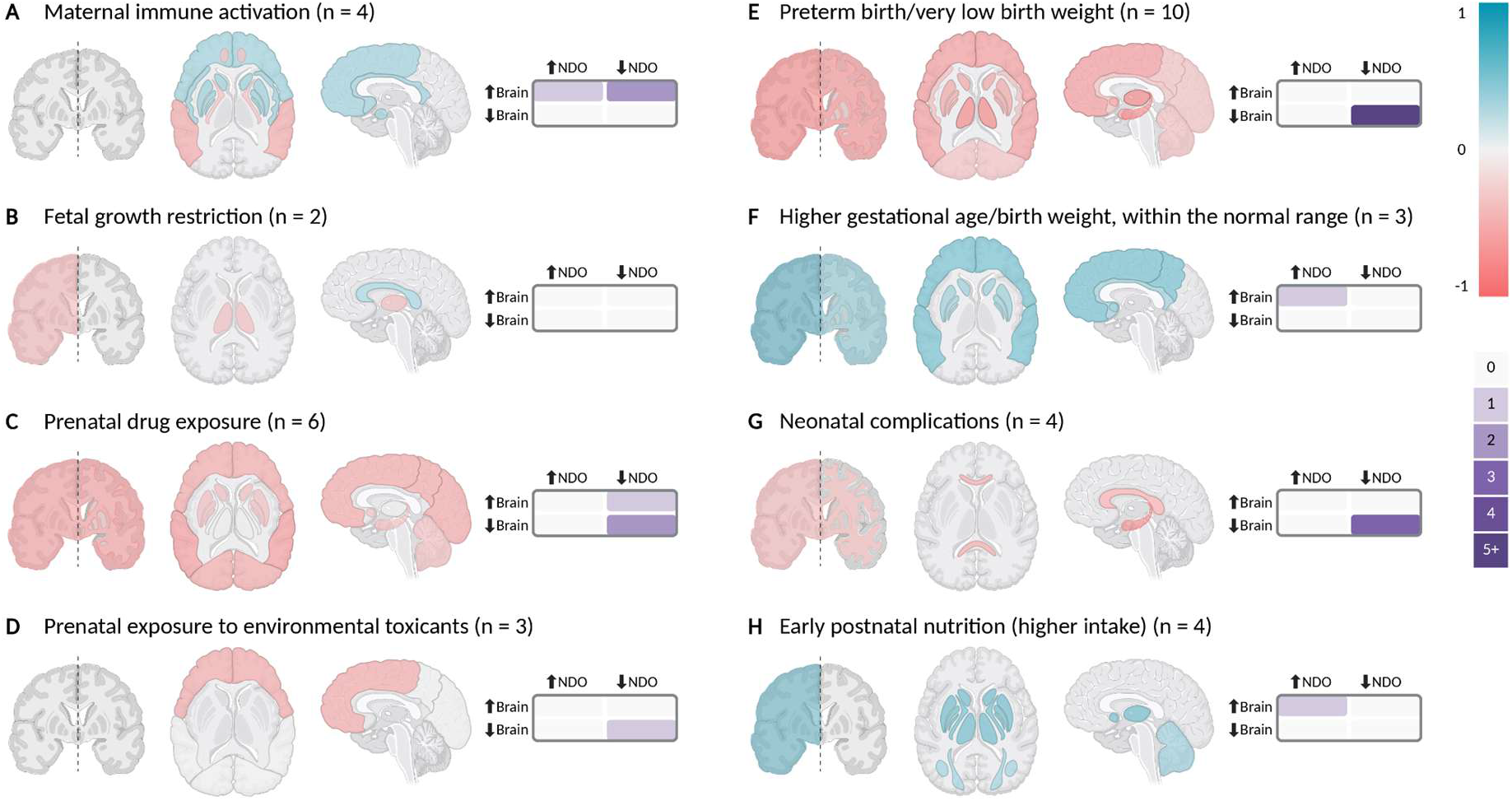
Overview of associations between different biological early-life exposures (**A**-**H**), the brain and neurodevelopmental outcomes. Associations with brain structure are shown separately for global (left brain) and regional changes (middle and right brain). Color indicates direction of association (blue = accelerated maturation, red = delayed maturation), and color intensity reflects consistency across studies, based on a Laplace-smoothed score from −1 to +1. The 2×2 grids show whether structural brain changes were associated with positive or negative neurodevelopmental outcomes, with color intensity encoding how frequently each combination occurred. NDO, neurodevelopmental outcomes. This figure was created in BioRender, https://BioRender.com.

*Fetal growth restriction* (n = 2; Figure 5, **B**). Fetal growth restriction was associated with lower total brain volume, altered cerebrospinal fluid distribution, and mixed microstructural integrity (45,67). It was linked to poorer motor development by 1 year (45,67), though mediation was not tested.

*Drugs* (n = 6; Figure 5, **C**). Prenatal drug exposure was consistently associated with widespread macrostructural reductions in volume, surface area, and cortical thickness. Alcohol exposure was linked to lower total brain volume, gray matter volume, and cerebellar volume, as well as reduced cortical thickness and surface area (68–70), with middle temporal gyrus measures mediating links to poorer cognition, executive functioning, and numerical skills. Continued tobacco exposure was associated with lower volumes and cortical thinning, with frontal thinning mediating higher affective problems in early childhood (71). Notably, offspring of mothers who stopped smoking during pregnancy showed no differences from the control group, suggesting a duration-dependent effect. Heroin and/or cocaine exposure was associated with higher hippocampal volumes and poorer memory at 14 years (72). Microstructural evidence was limited to one study showing accelerated maturation in the internal capsule and thalamus after tobacco-only exposure (73).

*Environmental toxicants* (n = 3; Figure 5, **D**). Higher prenatal air pollution exposure was associated with reduced frontal and temporal cortical thickness, which mediated a link to poorer inhibitory control in middle childhood (74). Chlorpyrifos exposure was similarly linked to reduced frontal and parietal cortical thickness, as well as bidirectional cortical surface changes and disrupted sexual dimorphism (75). In contrast, PFAS exposure showed no association with white matter microstructure but was linked to lower cognitive and language abilities (76).

##### Perinatal and postnatal

*Preterm birth/very low birth weight* (n = 10; Figure 5, **E**). At the macrostructural level, preterm birth and/or VLBW was associated with lower global and regional gray and white matter volumes, enlarged ventricles, and lower growth rates from infancy through early adulthood (45,61,77–81). Cortical thickness was higher at term-equivalent age but lower by middle childhood, and surface area was reduced with accelerated decline through childhood (78,82). These changes were associated with lower cognition, learning, executive functioning, language, and motor development from infancy through early adulthood (45,61,77–79,82). Microstructurally, lower integrity in central white matter tracts and the thalamus related to lower motor and cognitive development (45), and lower myelin water fraction mediated a link to lower executive functioning in early adulthood (83).

*Gestational age/birth weight within the normal range* (n = 3; Figure 5, **F**). Higher birth weight was consistently associated with increased global and regional gray and white matter volumes and greater surface area from early childhood through early adulthood (84–86), with larger anterior cingulate surface area relating to better reaction times. Combined lower gestational age and birth weight was associated with smaller caudate volumes and slower, more variable reaction times, without mediation testing (84).

*Neonatal complications* (n = 4; Figure 5, **G**). Complications associated with preterm birth, including parenteral nutrition, critical illness, mechanical ventilation, and exposure to the sedative midazolam, were associated with lower total brain volume, reduced hippocampal growth, and decreased white and gray matter integrity from the neonatal period through middle childhood, in turn relating to poorer cognitive and motor development (45,47,87). Delayed maturation caused by sedatives appeared to be specific to midazolam rather than fentanyl or morphine (87). Neonatal inflammation in a predominantly healthy term-born sample, however, showed no association with brain structure or behavior (46).

*Early postnatal nutrition* (n = 4; Figure 5, **H**). In preterm infants, higher caloric, protein, and lipid intake in the first weeks was associated with larger total brain, cerebellar, basal ganglia, and thalamic volumes and growth, with total brain and subcortical growth mediating better psychomotor development by 2 years corrected age (88,89). Higher lipid intake was also linked to higher white matter integrity in several tracts (88,89). Longer parenteral nutrition was associated with lower volumes and reduced microstructural integrity, likely reflecting greater overall illness severity in infants requiring parenteral nutrition (88). One study, however, found no associations between early nutrition in preterm babies, brain development, and neurodevelopmental outcomes (90). In term-born infants, one study found formula feeding associated with initially higher but later lower myelin water fraction, and with lower cognitive and verbal development scores compared to breastfeeding – however, effect sizes were small, all outcomes fell within the normal range, and mediation was not tested (91).

### Psychological Exposures

*Prenatal: maternal mental health* (n = 9; Figure 6, **A**). Prenatal stress was associated with lower fetal and infant hippocampal volumes, which mediated links to poorer socioemotional development and toddler cognition (92,93). Sex and timing modulated gyrification, with early prenatal stress linked to higher gyrification in women and lower in men in early adulthood, and stress in late pregnancy to lower gyrification in both sexes (62). Weather-related stress (hurricane exposure during pregnancy) was associated with higher amygdala and lower medial orbitofrontal volumes, and more externalizing behavior, without mediation testing (60). Prenatal depressive symptoms were associated with accelerated brain aging that mediated later anxiety and mood dysregulation (94), and with both accelerated and delayed maturation of brain microstructure (higher fiber density in the uncinate fasciculus at birth; lower integrity in the amygdala-frontal pathway and cingulum at 4 years). Microstructural change in both directions was related to greater socioemotional and externalizing difficulties, partly in a sex-specific manner (95,96). Prenatal anxiety was associated with both higher and lower microstructural integrity across regions, with no link to socioemotional development at 1 year (97). Poorer maternal sleep quality was associated with higher uncinate fasciculus integrity that mediated greater infant negative emotionality (98).

**Figure 6:**
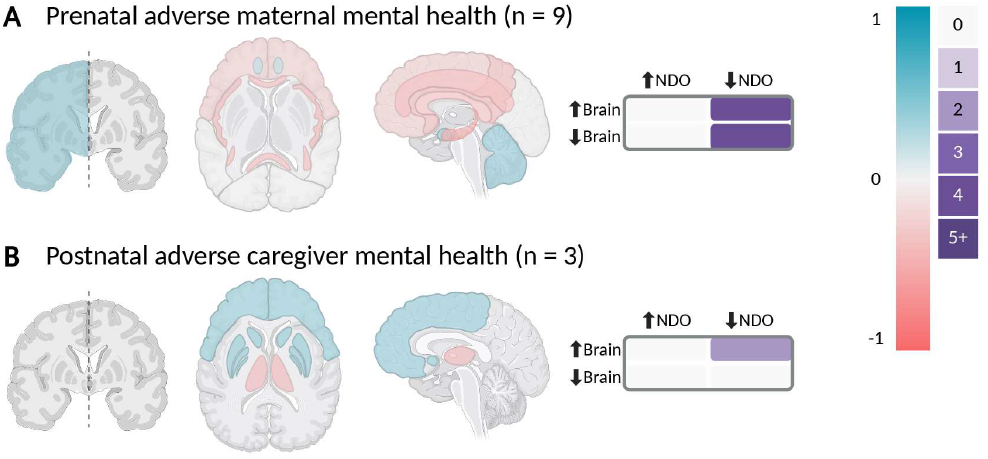
Overview of associations between different psychological early-life exposures (**A**, **B**), the brain and neurodevelopmental outcomes. Associations with brain structure are shown separately for global (left brain) and regional changes (middle and right brain). Color indicates direction of association (blue = accelerated maturation, red = delayed maturation), and color intensity reflects consistency across studies, based on a Laplace-smoothed score from −1 to +1. The 2×2 grids show whether structural brain changes were associated with positive or negative neurodevelopmental outcomes, with color intensity encoding how frequently each combination occurred. NDO, neurodevelopmental outcomes. This figure was created in BioRender, https://BioRender.com.

*Postnatal: caregiver mental health* (n = 3; Figure 6, **B**). A negative emotional caregiving environment was associated with smaller left hippocampal volumes and poorer episodic memory in adolescents with prenatal drug exposure, with the exposure-brain association reversing in boys by 18 years (99). Higher postnatal maternal anxiety and caregiver stress were mostly linked to accelerated maturation (higher pallidum volumes, higher frontal gyrification), which related to poorer emotion regulation, partly in a sex-specific manner (62,100).

*Screen time* (n = 2; no visualization). Higher infant screen time was associated with altered network integration interpreted as accelerated network maturation, though the two studies examined different network pairs with effects that are not easily reconciled (101,102). These topological changes mediated links to poorer emotion regulation, lower resilience, longer decision latency, and higher anxiety.

### Social Exposures

*Prenatal: sociodemographic factors* (n = 2; Figure 7, **A**). Lower prenatal SES was associated with both higher and lower regional volumes shortly after birth, and with poorer language development and fewer attention deficit hyperactivity disorder and oppositional symptoms at 2 years, though these outcomes were not linked to the brain alterations (103). Higher social support showed associations with bidirectional volume changes as well (103). Prenatal social disadvantage was linked to more externalizing and dysregulating symptoms at 2 years; these symptoms were associated with reduced global gray and white matter volumes, but direct exposure-brain associations and mediation were not tested (104).

**Figure 7:**
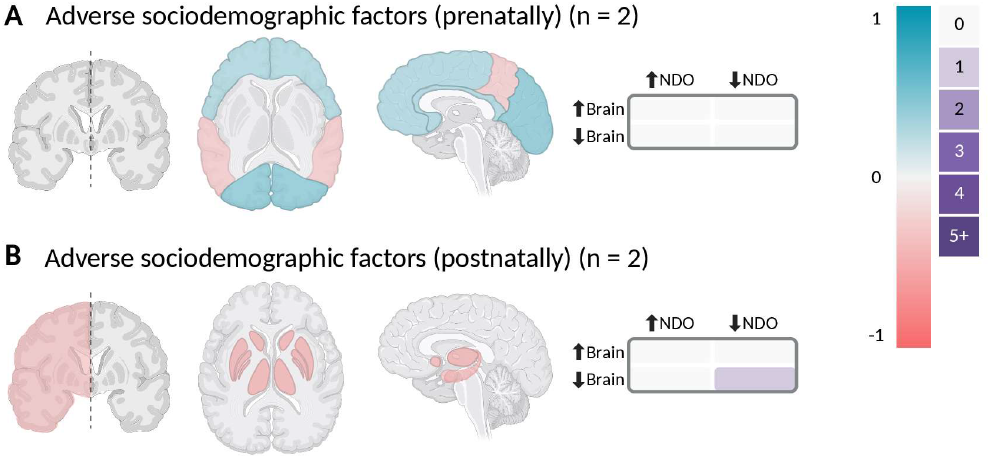
Overview of associations between different social early-life exposures (**A**, **B**), the brain and neurodevelopmental outcomes. Associations with brain structure are shown separately for global (left brain) and regional changes (middle and right brain). Color indicates direction of association (blue = accelerated maturation, red = delayed maturation), and color intensity reflects consistency across studies, based on a Laplace-smoothed score from −1 to +1. The 2×2 grids show whether structural brain changes were associated with positive or negative neurodevelopmental outcomes, with color intensity encoding how frequently each combination occurred. NDO, neurodevelopmental outcomes. This figure was created in BioRender, https://BioRender.com.

*Postnatal: sociodemographic factors* (n = 2; Figure 7, **B**). Lower maternal education and family income were predominantly associated with lower regional volumes from birth through early childhood, with lower pallidum and caudate volumes mediating the link to poorer visual reception (61). In preterm infants, greater socioeconomic deprivation was linked to higher psychopathology and poorer psychomotor development; these outcomes were related to lower global and regional volumes, but direct exposure-brain associations and mediation were not tested (105).

## Discussion

No period of life involves more rapid changes in the brain than the prenatal and early postnatal window, which makes it especially vulnerable to a wide range of exposures (1,2,7–9). This review asked how such exposures shape brain development, whether the resulting alterations relate to neurodevelopmental outcomes, and what patterns emerge across exposures. We found that diverse early-life exposures affect brain structure at both micro- and macrostructural levels, with these changes frequently associated with cognitive, motor, and socioemotional differences. Biological exposures tended to act through relatively direct mechanisms, whereas psychological exposures combined regionally distinct delayed and accelerated development within a single exposure. The field is growing, with psychosocial exposures only beginning to receive due attention.

### Biological exposures

For *maternal immune activation*, the bidirectional effects on brain maturation, particularly at the microstructural level, can be explained in the context of the diverse mechanisms through which maternal inflammation might affect the brain (7,38). Despite this bidirectionality, neurodevelopmental outcomes were mostly adverse.

*Fetal growth restriction* aligned with the broader literature documenting volume reductions and altered connectivity (39,106).

*Prenatal drug exposure* produced widespread alterations beyond dopamine circuits, largely indicating delayed maturation, in line with the broader literature (7,17,35), and with poorer outcomes persisting into adulthood, mediated by brain alterations in a subset.

*Exposure to environmental toxicants*, specifically to air pollution and chlorpyrifos, was associated with delayed maturation, particularly in frontal regions, that mediated the link to poorer neurodevelopmental outcomes when tested, consistent with the broader literature (22,41,107,108).

*Preterm birth and very low birth weight* showed a highly consistent adverse pattern, with mediation supported where tested, a finding that is consistent with the broader literature (19,32,109,110).

For *gestational age and birth weight within the normal range*, higher values were associated with accelerated macrostructural maturation and, in a subset, favorable outcomes, consistent with the small existing literature (111).

*Neonatal complications* were each covered by only one or two studies, yet severe complications, even when not primarily neurological, were consistently associated with delayed brain maturation and poorer outcomes; a single term-born study of subclinical inflammation found no effects, though this is preliminary.

*Early postnatal nutrition*, assessed mainly in preterm infants, showed higher macronutrient intake (lipids in particular) associated with greater brain volume and white matter integrity, and with better psychomotor development, which is supported by the broader literature (10,112). Breastfeeding was linked to higher myelination and better cognition.

### Psychological exposures

*Prenatal maternal mental health*, mostly assessed subclinically, produced a heterogeneous picture, with reduced hippocampal and enlarged amygdala volumes the most robust findings, alongside mixed alterations in gyrification, accelerated brain aging, and bidirectional microstructural change, which is also reflected in the broader literature (30,113). The limbic and prefrontal focus aligns with the high glucocorticoid receptor density in these regions, rendering them particularly susceptible to HPA-axis activations (7); it is further consistent with the poorer socioemotional outcomes, mediation-supported in several studies. Sex differences were rarely examined and mixed when reported, tentatively suggesting a differential vulnerability to adverse prenatal conditions (13).

For *postnatal caregiver mental health*, findings were heterogeneous, with reduced hippocampal volume but increased pallidum volume and frontal gyrification, the latter linked to poorer emotion regulation.

*Infant screen time* was assessed by only two studies from a single group examining network topology, both showing altered topology and poorer outcomes. This broadly aligns with the emerging literature reporting lower white matter integrity and reduced cortical thickness, with the prefrontal cortex appearing particularly vulnerable (49,114). Increased screen time has largely been associated with negative cognitive and psychosocial outcomes (20,115).

### Social exposures

*Adverse prenatal sociodemographic factors* were covered by two studies with bidirectional changes in volume. The broader, still limited, literature associates these factors predominantly with reduced volumes, though with some inconsistency (50,116). While social disadvantage was associated with poorer neurodevelopmental outcomes, the included evidence is insufficient to establish a mediating pathway via brain structure.

*Adverse postnatal sociodemographic factors* were mostly associated with lower brain volumes that in turn mediated poorer outcomes, though this is based on only two studies, one of which was a large multicohort study (61). The results align with the broader literature linking lower postnatal SES to reduced volumes and smaller surface area, particularly in language and executive-function regions (15,117).

### Overarching patterns

#### Biological versus psychological exposures

Biological exposures tended to act in a consistent direction per exposure, potentially reflecting more direct mechanistic pathways, whereas psychological exposures produced both accelerated and delayed maturation within a single exposure; their effects on neurodevelopmental outcomes nonetheless converged. Several non-mutually exclusive mechanisms may explain this. The most established is that adverse exposures stress the developing system, delaying or preventing normal development; this is largely supported for biological exposures and partly by the literature on psychosocial exposures. Alternatively, the stress acceleration hypothesis (118) predicts that adversity accelerates maturation as an adaptive response to environmental demands, potentially at long-term cost, a hypothesis that was also supported by some of the reviewed studies of psychosocial exposures. A third hypothesis is that adversity may produce genuinely adaptive change, leading to brain alterations that confer long-term resilience (119). This hypothesis had limited support from reviewed studies of maternal immune activation. These interpretations warrant caution. The heterogeneity observed for psychological exposures may partly reflect variation in how these factors were assessed and the grouping of distinct aspects of mental health, though bidirectional effects within individual studies suggest this cannot fully account for it. Subclinical effects may also be too small to produce consistent findings in smaller studies. Small to medium effect sizes have been reported (94,96), though replicability of these findings has not been assessed (120). Taken together, these findings caution against interpreting structural brain changes in isolation: neither accelerated nor delayed development straightforwardly maps onto better or worse outcomes, underscoring the need to examine all three domains together, ideally through mediation analyses.

#### Micro- versus macrostructure

When both were assessed, microstructure and macrostructure generally aligned in direction. A notable exception was prenatal maternal mental health, where the two dissociated, possibly reflecting the multiple etiologies of maternal mental health problems and/or differential sensitivity of these measures to psychological stressors.

### Trends and future directions

#### Current trends

Research into early-life exposures and their influence on neurodevelopment has gained momentum since the early 2010s, initially focused on biological exposures, with psychological and social exposures receiving attention mainly from 2015 onwards. The most studied exposures were prematurity and prenatal drug exposure. Many exposures received little attention, e.g., fetal growth restriction, individual aspects of pre- and postnatal parental mental health, infant screen time, and pre- and postnatal sociodemographic factors; exposures investigated only in single studies included parental age, maternal malnutrition, timing of cord clamping, and mother-child attachment security. Methods have evolved in parallel, from structural volumetrics and standard diffusion tensor imaging metrics toward more advanced diffusion approaches (neurite orientation dispersion and density imaging, fixel-based analysis), brain age, network topology, and multi-modal designs. The field has further moved from small group comparisons toward longitudinal growth curve modeling, mediation analyses, and large multi-cohort designs drawing on datasets such as the HEALthy Brain and Child Development (HBCD) study, the Adolescent Brain Cognitive Development (ABCD) study, the developing Human Connectome Project (dHCP), and the Baby Connectome Project (BCP), reflecting a recognition that individual research groups rarely recruit adequate samples.

#### Future directions

There is substantial scope for further work, particularly on the understudied psychological and social exposures. Parental mental health needs more differentiated investigation, potentially extending beyond subclinical presentations to clinically relevant distress and investigating protective factors as well. Beyond maternal mental health, the literature should broaden to include paternal mental health, since the rationale for focusing on the mother during pregnancy does not extend to the postnatal period (121). Infant screen time is a new and rapidly growing exposure warranting research to inform guidelines (49). For social exposures, SES’s influence on brain MRI measures has only recently become a research question in its own right (122), with its influence already well established in adolescent and adult samples (123–125) but scarce for the developing brain, particularly prenatally. Biological exposures are better studied, yet maternal and term-born postnatal nutrition remain underexamined within the full exposure-brain-outcome framework, as well as general timing and dosage effects. Methodologically, large multi-site datasets like HBCD, dHCP, or BCP will help to address persistent sample-size limitations. Longitudinal designs with higher temporal resolution would help elucidate timing effects and disentangle prenatal from postnatal exposures. Examining the exposure, the brain, and the outcome within the same population, ideally through mediation, remains crucial, since partial pathways risk being misleading. A central challenge is disentangling the many correlated biopsychosocial exposures acting on the developing brain (72), which will require both advanced statistics and rich, large-scale datasets.

### Strengths and limitations

This review synthesizes roughly 25 years of research and is holistic in two senses: it includes only studies assessing the full exposure-brain-outcome pathway within the same population, and it spans the full range of biopsychosocial early-life exposures, allowing cross-cutting patterns to emerge. Although individual studies are often small, the combined sample lends reasonable weight.

Several limitations apply. First, genetic influences were not considered, despite likely gene-environment interactions (5), so the associations reported here should be understood as reflecting environmental influences within a context that genetics also shape. Second, not all included studies analyzed every pairwise association, and only a subset conducted formal mediation, so confidence in the full pathway varies. Third, requiring all three elements of the exposure-brain-outcome pathway reduced the studies available per exposure. Therefore, we contextualized findings against the wider literature where possible. It also means the observed increased emphasis on psychosocial influences cannot be firmly established as a field-wide trend, though it is striking enough to likely reflect one. Fourth, the review assumes a unidirectional exposure-to-brain-to-outcome pathway, whereas behavior can also shape brain development (60), so reverse or reciprocal effects cannot be ruled out. Finally, imaging timepoints varied substantially, from the fetal period to late adulthood; while this limits direct comparisons, it also reveals how early-life exposures may have lasting associations across the lifespan.

## Conclusion

Early-life exposures are a promising area for future research, particularly the influence of psychosocial exposures on brain development and neurodevelopmental outcomes. This review showed that a range of early-life exposures shape brain development and, in turn, neurodevelopmental outcomes. Yet the mechanisms, how these effects unfold across the lifespan, and how exposures interact remain largely unclear. Future work should prioritize large, longitudinal datasets with rich information about the developing child and their environment to understand these pathways – knowledge essential for providing the best possible environment for a child to grow.

## Acknowledgments & Disclosures

J.D. was supported by a Gates Cambridge Scholarship (#OPP1144); H.B. was supported by a scholarship of the G. C. Grindley Fund; A.A.B. was supported by BIMH R01MH133843, R01MH132934, R01MH134896; R.A.I.B. was supported by an Academy of Medical Sciences Springboard Award and the HDRUK Molecular to Health Records program.

R.A.I.B is a co-founder of and holds equity in Centile Bioscience Inc. A.A.B. has equity in, consulted for, and has an inventorship interest in intellectual property licensed to Centile Bioscience Inc. All other authors report no biomedical financial interests or potential conflicts of interest.

